# Anthrax toxin receptor 2 is the Receptor for *Clostridium perfringens* NetF: Structural Insights into Toxin Binding and Pore Formation

**DOI:** 10.1101/2025.07.02.662835

**Authors:** Chang Wang, Filippo Cattalani, Ioan Iacovache, Arunasalam Naguleswaran, Faezeh Farhoosh, Jan Franzen, Laurence Abrami, F. Gisou van der Goot, Horst Posthaus, Benoît Zuber

## Abstract

Hemolysin β-pore-forming toxins (βPFTs) are key virulence factors of *Clostridium perfringens*, associated with severe diseases in humans and animals. Yet, the mechanisms by which *Clostridium* βPFTs recognize and engage specific target cells remain poorly understood. Here, we identify the cellular receptor for *C. perfringens* necrotizing enteritis toxin F (NetF), a recently discovered toxin implicated in severe enteritis in dogs and foals. We show that NetF binds to the same receptor as anthrax toxin, namely ANTXR2. Using cryo-electron microscopy, we determined the structure of the oligomeric NetF pre-pore as well as the transmembrane pore, both alone and in complex with the extracellular domain of ANTXR2. Unlike anthrax toxin, which binds to the apical MIDAS motif of ANTXR2 – as does the natural ANTXR2 ligand collagen type VI – NetF engages the receptor laterally, spanning both the von Willebrand A and the Ig-like domains. This interaction positions the toxin near the membrane, facilitating contact with membrane lipids and promoting transmembrane pore formation. Our findings uncover key principles of hemolysin βPFT-receptor recognition and advance our understanding of how pathogenic bacteria use these toxins to breach host defenses.

## INTRODUCTION

*Clostridium perfringens* is a versatile pathogen, causing wound infections, septicemia, enterotoxemia or enteritis in animals and humans ^1–3^. Especially, intestinal diseases in animals such as pigs, cattle, sheep, goats, chickens, dogs and horses result in significant losses of livestock and companion animals worldwide. *C. perfringens* induces tissue damage via highly potent exotoxins of which hemolysin β-pore-forming toxins (βPFTs) represent the largest group, with currently 11 identified members^4,5^. Two of these, *C. perfringens* beta-toxin (CPB) and necrotic enteritis toxin B (NetB), are essential virulence factors in necrotic enteritis in pigs and humans and poultry, respectively. Our knowledge about other *C. perfringens* hemolysin βPFTs, however, is limited. In 2015, a novel hemolysin βPFT called NetF was identified in *C. perfringens* isolates from dogs with hemorrhagic gastroenteritis and foals with necrotic enterocolitis ^6^. Subsequent epidemiological evidence substantiated, that the toxin plays an important role in *C. perfringens* induced enteric disease in these animals ^7–11^. NetF molecular weight is 34.4 kDa, it shares 30%, 33% and 48% sequence identity with *S. aureus* α-hemolysin (Hla), CPB and NetB, respectively. It was shown to bind to the membrane of susceptible cells through interaction with an undefined sialylated receptor and to form oligomers with varying stoichiometries on liposomes ^12^.

Hemolysin βPFT are mostly secreted by the bacteria as soluble monomers. They then bind to receptors on target cells, oligomerize into pre-pores and subsequently undergo conformational change to insert into the plasma membrane as ion permissive pores ^13^. Thus, identifying membrane receptors and the structural basis of the interaction between the toxin, the receptor and the plasma membrane is key to understanding the function of these toxins. So far, only two receptors for clostridial hemolysin βPFTs have been identified: CD31 for *C. perfringens* beta-toxin (CPB) and ganglioside GM2 for delta-toxin ^14,15^. In addition, oligomeric pore-structures have only been determined for CPB and NetB ^14,16,17^, whereas precise structural information about clostridial toxin-receptor and toxin-membrane interaction is still lacking.

Here, we elucidate the structure of NetF in its oligomeric pre-pore and its membrane-inserted pore state. We identify anthrax toxin receptor 2 (ANTXR2), also known as capillary morphogenesis gene 2 (CMG2), as a receptor for NetF and show that the toxin can efficiently target human, bovine, equine and canine ANTXR2. In addition, we determine the co-structure of NetF oligomer in complex with the extracellular part of ANTXR2, revealing novel insight into the interaction between a clostridial hemolysin βPFT and its membrane receptor.

## RESULTS

### Structural characterization of NetF

Previous research demonstrated that NetF oligomerizes in different stoichiometries, forming complexes composed of six to nine protomers ^12^. We recombinantly expressed the toxin in *Escherichia coli* and confirmed that NetF binds and oligomerizes on synthetic membranes. Cholesterol increased membrane binding of NetF (Sup. Fig. 1AB). To optimize the conditions for cryo-EM characterization of NetF oligomeric form, we generated pores following two distinct methods: 1) incubation of NetF monomers with liposomes, leading to the formation of oligomeric pores, which were subsequently solubilized (Sup. Fig. 1C), and 2) incubation of NetF monomers with pre-formed lipid nanodiscs (Sup. Fig. 1D). Surprisingly, the latter led predominantly to nonameric pre-pores (Figure 1AB), while detergent-solubilized oligomers mostly adopted an octameric pore conformation (Figure 1CD), as monitored by cryo-EM. The structures of NetF pre-pore and pore represent a hemolysin-like oligomer organized into three distinct regions: the cap, the rim and the stem (Figure 1CE). The high resolution cryo-EM maps enable precise atomic modeling of the backbone and most sidechains, except for the first eleven N-terminal amino acids, which are partially unstructured (Sup. Fig. 2).

**Figure 1.**
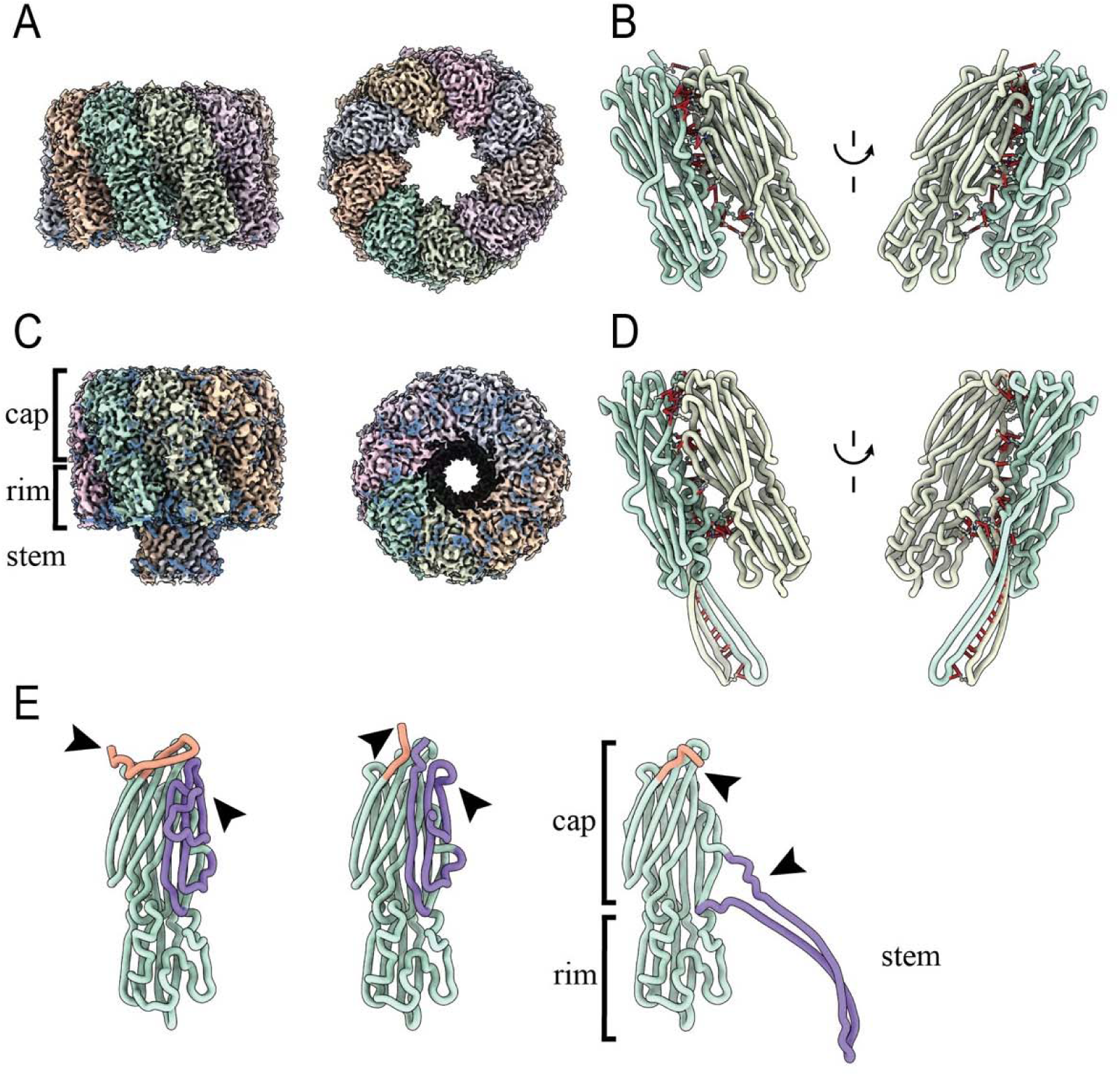
Structure of NetF oligomers and structural changes required for pore formation. **(A)** Cryo-EM map of NetF on MSP2N2 lipid nanodiscs containing DOPC:DOPG:Cholesterol. Upon NetF addition nine monomers bind and oligomerize into a pre-pore conformation. Consecutive protomers are colored by a pastel rainbow. Extra densities are shown in blue. **(B)** The structures of two consecutive protomers extracted from the pre-pore are shown in cartoon representation and colored as in A. Residues involved in protomer – protomer interaction are shown as ball and stick with the distances shown as red dashed lines. Two representations rotated by 180° are shown for clarity. **(C)** Cryo-EM map of NetF pores after oligomerization on DOPC:DOPG:Cholesterol liposomes and solubilization in DDM:CHS. Eight NetF monomers bind and oligomerize to form the membrane pore. Consecutive protomers in the oligomer are colored as in A with extra densities shown in blue. The position of the cap, rim and stem is shown. **(D)** The structure of a protomer extracted from the pore is shown in cartoon representation as in (B). Residues involved in inter-protomer interactions are shown in ball and stick representation with the inter-residue distance shown as red dashed lines. Two views are shown for clarity rotated by 180°. **(E)** Comparisons between the soluble NetF monomer (AlphaFold prediction) on the left, a protomer extracted from the NetF pre-pore (middle) and a protomer extracted from the pore (right). The position of the N-terminus is shown in orange and emphasized by a black arrow (top). While the monomer prediction shows the N-terminus part of a β-sheet of the cap domain (left) it moves upward from its position in the pre-pore (middle). This movement is accompanied by the pre-stem shifting towards the position previously occupied by the N-terminus (purple, emphasized by lower black arrow). Upon membrane insertion the pre-stem flips towards the membrane (purple and marked by the lower arrow). The N-terminus flips down to the position previously occupied by the pre-stem loop into the entrance cavity of the pore.

In the absence of X-ray data, the NetF soluble monomeric structure was predicted using Alphafold3 ^18^ (Figure 1E, Sup. Fig. 3A). A comparison between the AlphaFold3 monomer prediction and a protomer extracted from the NetF pre-pore indicates that AlphaFold3 prediction is extremely accurate with a root-mean-square deviation (RMSD) of 1.9 Å across all amino acids. Although the overall prediction confidence score is high (pTM of 0.91), the N-terminal region, the pre-stem loop, and one loop of the rim domain display lower confidence (Sup. Fig. 3A). The differences between the predicted monomer and the protomer, as well as comparison with other hemolysin βPFTs, suggest that multiple conformational changes are necessary for oligomerization (Figure 1E). The oligomerization of NetF into pre-pore conformation involves extensive interactions (Table S1 and S2), engaging a large area on each side of the cap and rim domains, measuring approximately 1400 Å^2^ and encompassing more than 50 residues (Figure 1B, Sup. Fig. 2B). The N-terminus, however, remains flexible and unresolved (residues 1-20). In the pre-pore conformation, the pre-stem loop is partially visible except for five residues (Figure 1E). Notably, in contrast to the AlphaFold3 prediction of NetF monomer and to the monomer structure of other known hemolysins, the pre-stem of the pre-pore is moved upwards, away from its position on the cap domain. This structural rearrangement may play a critical role in pore formation, potentially priming the protein for membrane insertion.

The NetF pore structure reveals additional conformational changes required for membrane insertion. In the pre-pore, the pre-stem loop is folded against the cap domain, and its extraction during pore formation triggers a second rearrangement of the N-terminus. Although unresolved in both structures, the N-terminus position can be inferred from its orientation and diffuse density of the backbone and N-terminal His-tag (Figure 2AB). In the pre-pore, it points outward, but upon pre-stem loop unfolding, it shifts toward the cavity formed by the cap domain, loosely occupying the site of the pre-stem (Figure 2B). As the β-barrel forms, inter-protomer interactions expand by 50% to ∼2100 Å^2^ (Table S1 and S2), involving 90 residues (Figure 1B, Sup. Fig. 2D). Notably, the cap and rim domains remain largely unchanged during this transition.

**Figure 2.**
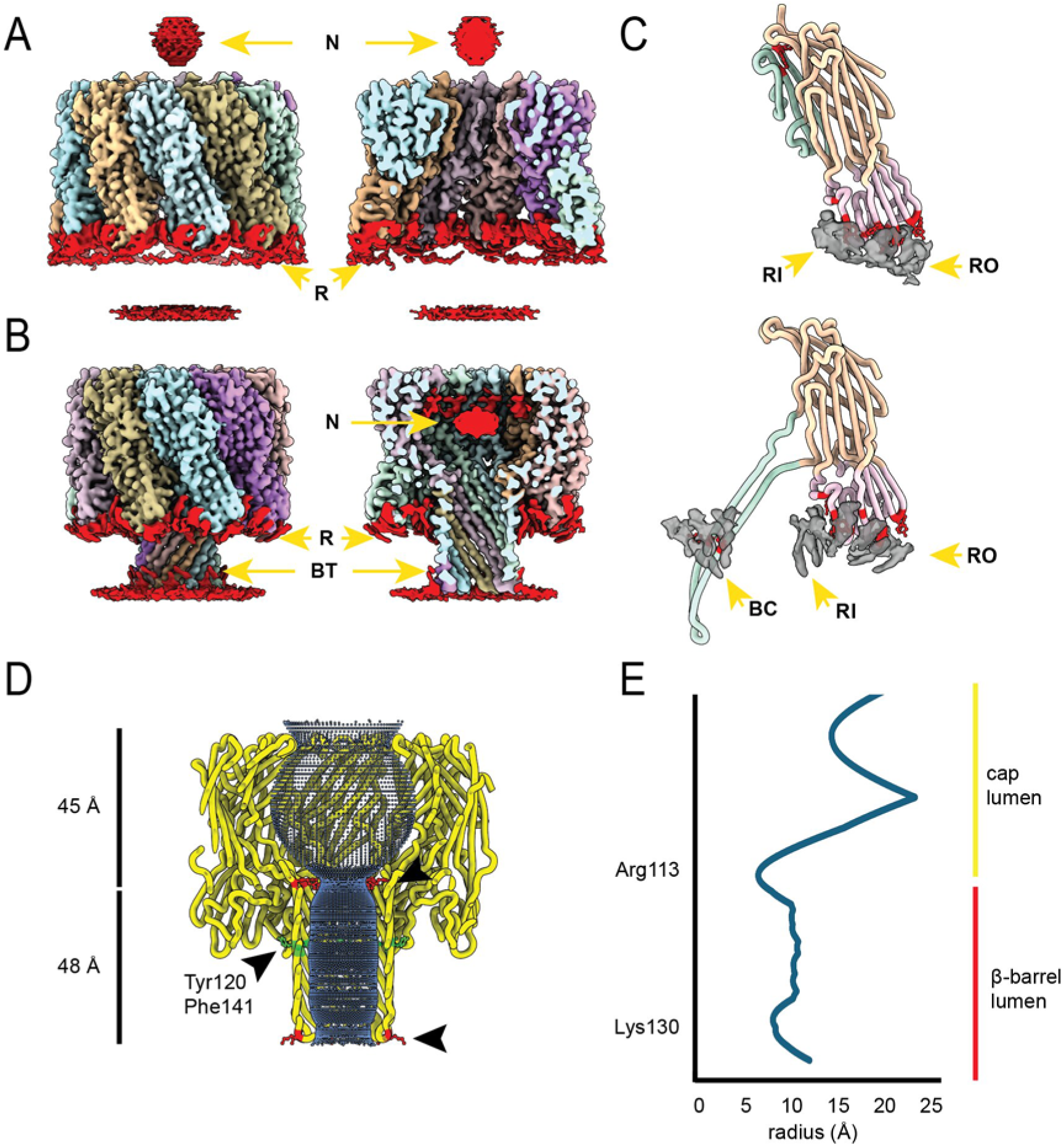
Characterization of NetF – lipid interaction and the transmembrane pore. **(A)** Cryo-EM map of the NetF pre-pore showing the extra densities (red) of the hexa-histidine tagged N-terminus (N) and lipid molecules bound to the rim domain (R). Side view of the pre-pore (left) shown with each protomer colored and extra densities in red. The density of the N-terminus (N) is seen above the oligomer while lipids bound to the outer rim domain are clearly visible. Median cut-through view of the pre-pore in side-view shows the empty cap cavity and lipid densities bound to the inner rim domain reveals lipid densities bound to the inner rim domain. **(B)** Cryo-EM map of the NetF pore (left) showing extra densities of lipids and detergents bound to the rim domain (R) and the β-barrel trans side (BT). A median cut-through view (middle) shows clearly the density of the hexa-histidine tagged N-terminus (N) now positioned in the middle of the cap cavity. Density from the N-terminus is seen on the inner cap domain. Lipid and detergent molecules (red) are seen interacting with the rim domain on the inner and outer surfaces of the rim (R). In addition, lipid molecules are seen interacting directly with the β-barrel on the trans cytoplasmic side and outer cis side. **(C)** Structure of a NetF protomer rim domain extracted from the pre-pore structure (top) is shown with the cap domain in orange, rim domain in pink and pre-stem loop in green. In gray lipid densities are shown bound to the rim domain on the outer surface (RO) and inner surface (RI). Aromatic residues in the rim and stem domains are highlighted in red and shown in ball-and-stick representation. Structure of a NetF protomer – rim and stem domains – extracted from the pore structure (bottom) is shown color coded as before. The densities of lipid molecules (gray) are shown on the outer rim (RO), inner rim (RI) and the aromatic ring (red) on the cis side of the transmembrane β-barrel (BC). Aromatic residues involved in the lipid interaction are highlighted in red and shown in ball-and-stick representation. **(D)** Highlight of the inner cavity of the NetF pore is shown in blue as calculated using HOLE. A large cavity on the cis side is formed by the oligomerization of the cap domains. Arginine 113 on the stem loop forming the transmembrane β-barrel forms the tightest constriction on the cis side. A second constriction on the trans/cytoplasmic side is formed by the β-turns flipping inwards constricting the barrel. The dimensions of the central cavity and the barrel cavity are shown on the left. **(E)** Plot of the radius of the NetF pore from **(D)** showing the two constriction sides formed by arginine 113 on the cis side and the β-turn at the position of lysine 130 on the trans side.

The most significant rearrangement during pore formation is the refolding of the pre-stem loop into a 50 Å-long transmembrane β-barrel, characterized by a hydrophilic lumen and a hydrophobic exterior. The barrel has a diameter of approximately 10 Å, with two constrictions that narrow the lumen to 6.7 Å on the cis (i.e. extracellular) side and 8.2 Å on the trans (i.e. cytoplasmic) side (Figure 2DE). The lumen of the transmembrane pore is polar but contains no charged residues, except for an arginine residue (Arg 113) at the cis entrance and two lysine residues (Lys 130 and 131) on the trans side, both of which contribute to the constriction sites. While the Lys 130 points away from the lumen it bends the β-turn inwards creating a second constriction. An aromatic belt formed by residues Tyr 120 and Phe 141 is positioned on the cis side oriented towards the lipids, consistent with the typical features of transmembrane β-barrels (Figure 2C-D).

### NetF interacts with lipids through the rim and stem domains

The cryo-EM maps also reveal substantial extra densities surrounding the rim and stem domains of NetF oligomer, in particular in the pore conformation (Figure 2A-C). Based on their shape and chemical environment, these densities correspond to bound lipid and detergent molecules. They are located around loops containing aromatic and charged residues: Tyr 74, Tyr 75 and Trp 254 bind lipids on the outer rim domain; Tyr 184, Trp 259, His 199 interact with lipids at the protomer-protomer interface. Furthermore, clear lipid/detergent densities are visible tightly interacting with the aromatic belt (Tyr 120 and Phe 141) as well as the middle and trans parts of the transmembrane β-barrel (Figure 2BC). It has been previously demonstrated for *S. aureus* α-hemolysin and related toxins that a loop in the rim domain interacts with lipid headgroups ^19,20^. The equivalent loop is however shortened in NetF and lacks the tryptophan residue required for lipid binding (Trp 179, Sup. Fig. 3B), as also observed in *C. perfringens* β-toxin and NetB ^17,21^. The presence of NetF-bound lipids and detergents suggest that the rim domain is partially embedded in the membrane tightly interacting with lipids and cholesterol.

### NetF resistance is associated with reduced ANTXR2 expression

We next set out to identify the cell surface receptor for NetF. To determine suitable cell lines, we performed viability assays on different human, murine, canine, equine and bovine cells. NetF was toxic to human U937 and HFF-1 cells, canine MDCK and glial J3T cells, primary equine fibroblasts and bovine endothelial BUcEC cells. In contrast, other cell lines from these species as well as two murine cell lines were resistant to NetF (Figure 3A).

**Figure 3.**
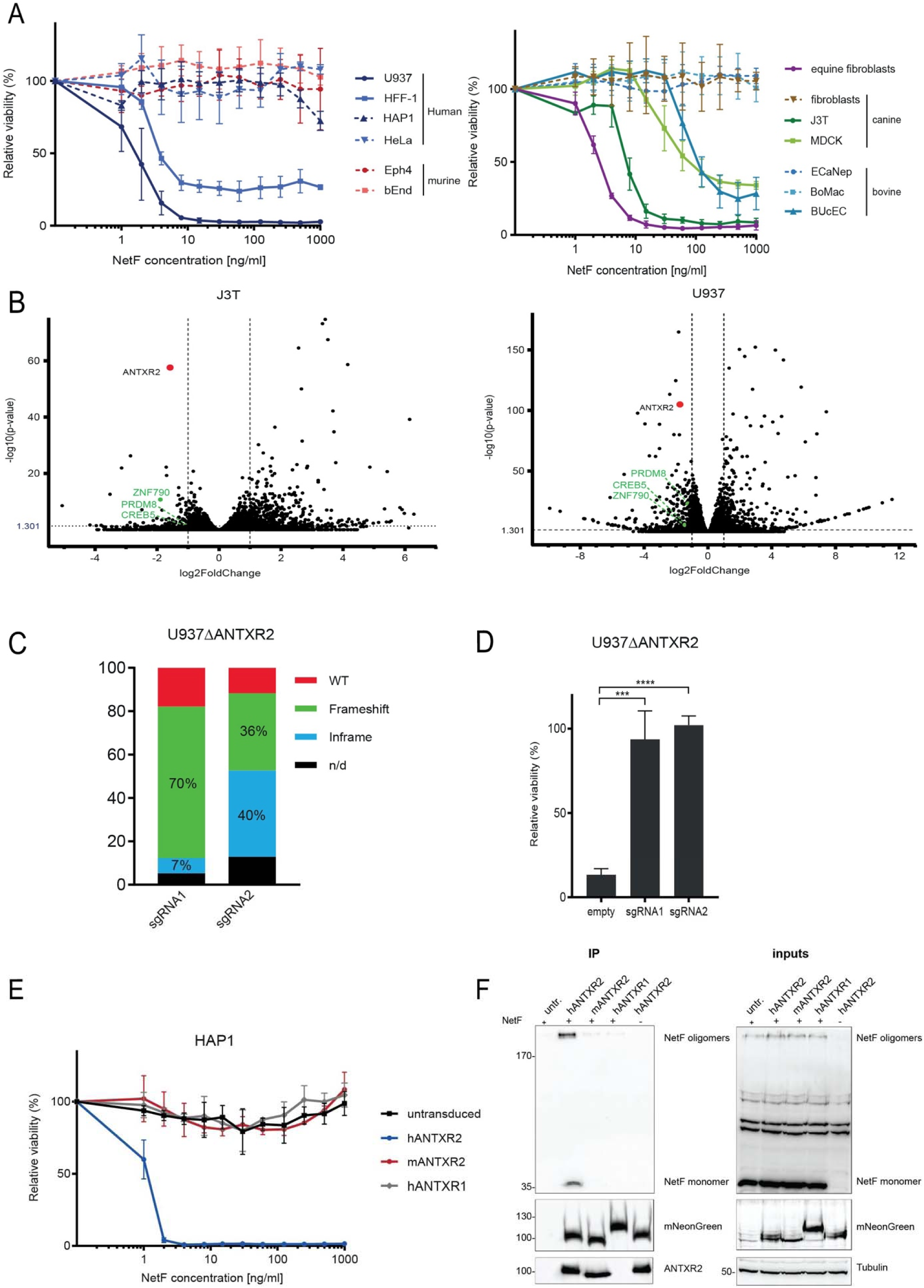
ANTXR2 interacts with NetF and is essential for NetF cytotoxicity. **(A)** Viability of human, murine, equine, canine and bovine cell lines after incubation with indicated doses of NetF (24 h, 37°C), in % to untreated control cells. Data are represented as means (n=9) ± SD. **(B)** Volcano plots depicting gene-expression profiles of NetF-resistant U937 and J3T cells compared to the respective NetF-sensitive wild type polyclonal cell population. ANTXR2 stood out as the most relevant down-regulated hit present in both experiments. **(C)** TIDE analysis showing high percentage of frameshift and in frame mutations introduced in the *ANTXR2* gene after sgRNA1- and sgRNA2-mediated ANTXR2 knockout in U937 cells. **(D)** Viability of U937 cells targeted by sgRNA1- and sgRNA2-mediated ANTXR2 knockout after incubation with 1 µg/ml NetF (24 h, 37°C), in % to untreated control cells. The empty lentiviral vector (empty) was used as control. Data are represented as means (n=9) ± SD. One-way ANOVA, Sidak’s multiple comparison test, **** (p < 0.0001), *** (p < 0.001). **(E)** Viability of HAP1 cells expressing mNeonGreen-tagged human ANTXR2, mouse ANTXR2 and human ANTXR1 after incubation with indicated NetF doses (24 h, 37°C), in % to untreated control cells. Data are represented as means (n=9) ± SD. **(F)** Westen blot analysis of co-IP experiments in HAP1 cells expressing human ANTXR2, mouse ANTXR2 and human ANTXR1, incubated with 10 µg/ml NetF. HAP1hANTXR2 cells without addition of NetF (-) were used as negative control. IP was performed using anti-mNeonGreen beads and showed enriched pull-down of NetF when coupled with hANTXR2. Immunoblot containing 50% eluate (IP) was performed on the same membrane with anti-His antibody to detect NetF and anti-mNeonGreen antibody (pulldown efficiency control). An additional immunoblot was performed with equal sample volumes and probed with anti-ANTXR2 antibody. Immunoblot containing 5% loading fraction (inputs) was performed on the same membrane using anti-His, anti-mNeonGreen and anti-tubulin (loading control) antibodies. Figure shows one of three replicates of the same experiment.

A small fraction of cells in several susceptible cell lines consistently survived NetF exposure, regardless of the toxin concentration applied. We chose the human U937 and the canine J3T cell lines, which showed less than 10% viability levels after 24 hours of toxin incubation (Figure 3A), to evaluate whether surviving cells were NetF-resistant subpopulations. Following six passages under constant toxin pressure, we isolated resistant subpopulations of both cell lines (U937-R and J3T-R) (Sup. Fig. 4A). Compared with the parental cell populations mRNA expression profiles of these resistant subclones showed four significantly downregulated genes in both U937-R and J3T-R cells: *ANTXR2, PRDM8, CREB5,* and *ZNF790* (Figure 3B and Sup. Fig. 4B). *ANTXR2* was the only gene encoding a transmembrane protein. *PRDM8,* encoding an intracellular histone methyltransferase, is a neighboring gene to *ANTXR2* and is co-regulated with it in the same transcriptional unit ^22^. *CREB5* encodes a transcription factor that binds to cis-responsive elements (CRE) in the promoter region of the *ANTXR2* transcriptional unit ^22^. No link has been identified between ANTXR2 expression and zinc finger protein 790 (ZNF790), a nuclear protein predicted to be involved in transcriptional regulation ^23,24^.

ANTXR2 is one of two receptors for anthrax toxin protective antigen (PA). The other PA receptor is ANTXR1, also known as tumor endothelial marker 8 (TEM8) ^25^. ANTXR1 and ANTXR2 are single pass transmembrane proteins with two extracellular domains: an N-terminal von Willebrand A (vWA) domain containing a metal ion-dependent adhesion site (MIDAS) motif and a membrane proximal immunoglobulin (Ig) domain ^26^. Additionally, they contain a C-terminal cytoplasmic domain involved in ligand-mediated signaling. ANTXR2 binds collagen VI via the MIDAS motif and participates in extracellular matrix homeostasis. Our RNA-sequencing (RNA-seq) data indicated that ANTXR1 expression was unchanged in U937-R and J3T-R compared to their susceptible counterparts (Dataset 1). These results led us to hypothesize that ANTXR2 serves as the membrane receptor for NetF.

### ANTXR2 interacts with NetF and is essential for its cytotoxicity

To validate the role of ANTXR2 in NetF toxicity, we performed CRISPR-Cas9-mediated knockouts (KOs) of ANTXR2 in U937 cells using two single-guide RNAs (sgRNAs) which led to a high number of frameshift and/or in-frame mutations (Figure 3C). Viability assays showed that sgRNA1- and sgRNA2-targeted cells became resistant to NetF treatment (Figure 3D) even though NetF binding and oligomerization were still detectable by western blot analysis on U937-R (Sup. Fig. 4C) as well as ANTXR2-KO U937 (Sup. Fig. 4D). PA binding was markedly reduced in U937sgRNA1 and U937sgRNA2 cells compared to cells treated with a control empty vector (Sup. Fig. 4E). This indicated successful depletion of ANTXR2 from the surface of U937 cells. These results were confirmed using HFF-1 cells where disruption of the ANTXR2 gene by either sgRNA1 or sgRNA2 led to a significant increase in cell viability upon NetF treatment (Sup. Fig. 4FG).

We next overexpressed mNeonGreen-tagged human ANTXR2 (hANTXR2), murine ANTXR2 (mANTXR2) and human ANTXR1 (hANTXR1) as a control in HAP1 cells. Expression, localization and functionality of the constructs were confirmed by western blot, PA binding assays (Sup. Fig. 5A) and immunofluorescence (IF) (Sup. Fig. 5B). Ectopic overexpression of hANTXR2 rendered HAP1 cells highly susceptible to NetF, unlike overexpression of hANTXR1 (Figure 3E). Furthermore, mANTXR2 did not confer susceptibility to NetF, consistent with our observation that all tested murine cell lines were resistant.

To assess interaction between ANTXR2 and NetF, we performed co-immunoprecipitations (co-IPs) from HAP1 cells overexpressing hANTXR2, mANTXR2, or hANTXR1 using anti-mNeonGreen-coated beads. We were able to co-precipitate NetF monomers and SDS-resistant oligomers in cells overexpressing human ANTXR2, while NetF co-precipitation was reduced in cells overexpressing mANTXR2 or hANTXR1 (Figure 3F).

### NetF targets ANTXR2 across different animal species

Since NetF was identified in canine and equine enteritis, we tested whether ANTXR2 ortholog expression sensitizes HAP1 cells to NetF. We ectopically expressed these orthologs as C-terminally mNeonGreen-tagged proteins (Figure 4A). Equine and bovine ANTXR2 localized correctly to the plasma membrane (Sup. Fig. 6A) and conferred susceptibility (Figure 4B), whereas canine ANTXR2 (cANTXR2) was largely retained intracellularly (Sup. Fig. 6A), preventing assessment. To overcome this, we expressed cANTXR2 in HEK293 cells alongside hANTXR2, mANTXR2, and hANTXR1 as controls (Figure 4C, Sup. Fig. 6A). Despite lower surface expression, cANTXR2 rendered HEK293 cells highly susceptible to NetF (EC50: 1-2 ng/ml) (Figure 4C), while mANTXR2-expressing cells were susceptible only at much higher NetF concentrations (EC50: 125 ng/ml). These results demonstrate that NetF toxicity depends on the surface expression of ANTXR2 and that it interacts with equine, bovine and canine ANTXR2.

**Figure 4.**
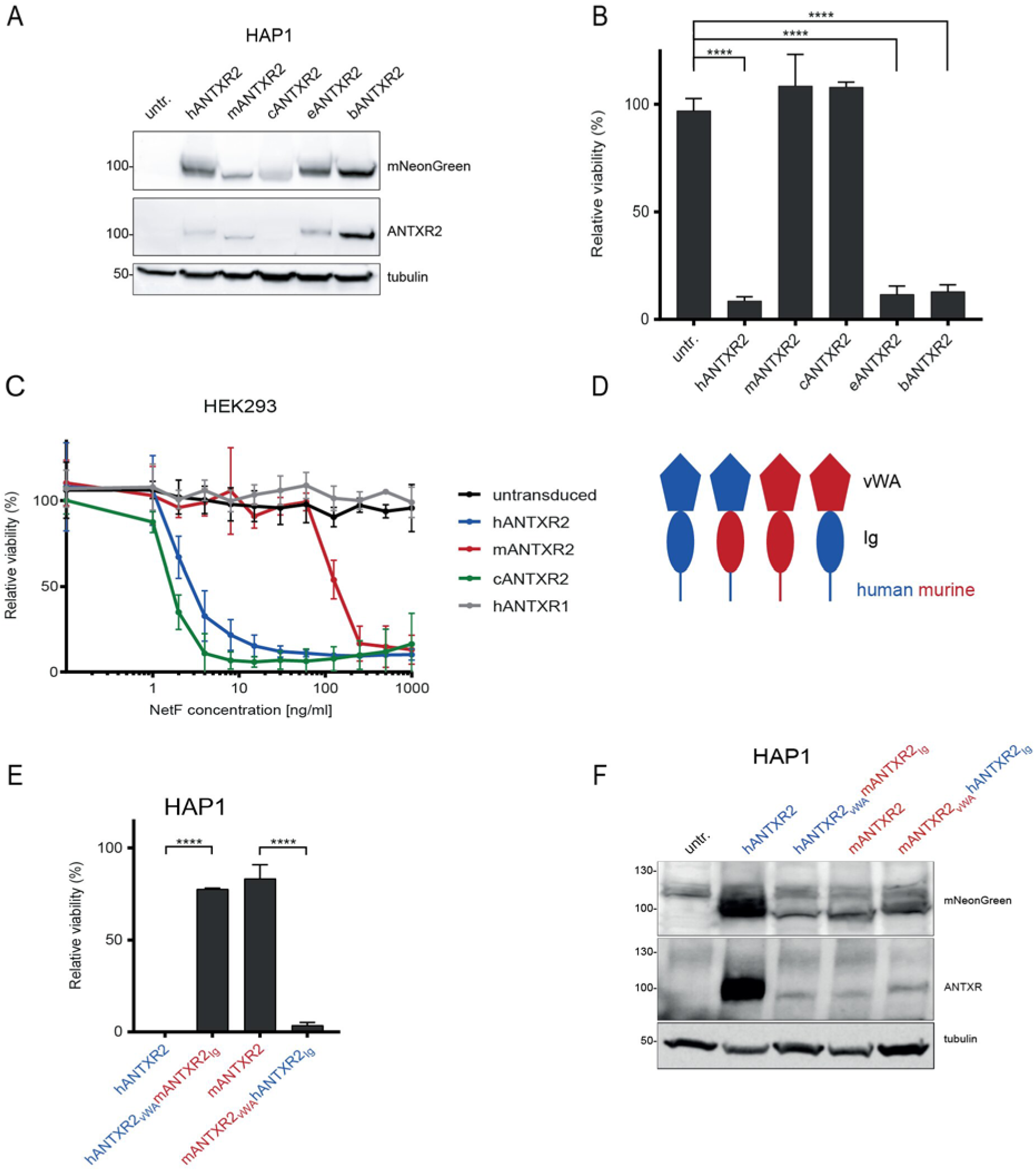
NetF interacts with ANTXR2 orthologs of many animal species. **(A)** Western blot analysis for expression levels of HAP1 cells transfected with various mammalian ANTXR2 orthologs. Immunoblots were probed on the same membrane with anti-mNeonGreen and anti-tubulin (loading control) antibodies. An additional western blot was performed with equal sample volumes and probed with anti-ANTXR2 antibody. **(B)** Viability of HAP1 cells expressing different mammalian ANTXR2 orthologs after incubation with 1 µg/ml NetF (24 h, 37°C), in % to untreated control cells. Data are represented as means (n=9) ± SD. One-way ANOVA,Sidak’s multiple comparison test, **** (p < 0.0001). **(C)** Viability of HEK293 cells expressing different mammalian ANTXR2 orthologs after incubation with 1 µg/ml NetF (24 h, 37°C), in % to untreated control cells. Data are represented as means (n=9) ± SD. **(D)** Schematic drawing of the extracellular and transmembrane regions of recombinant ANTXR2 WT and chimeric molecules used for ectopic expression in mammalian cells. Human: blue, mouse: red, vWA: von Willebrand A domain Ig: Ig domain **(E)** Viability of HAP1 cells expressing indicated chimeric constructs (Sup. Fig. 7AB) after incubation with 1 µg/ml NetF (24 h, 37°C), in % to untreated control cells. Data are represented as means (n=9) ± SD. One-way ANOVA, Sidak’s multiple comparison test, **** (p < 0.0001). **(F)** Western blot analysis for expression levels of HAP1 cells expressing human-mouse ANTXR2 chimeric constructs. Immunoblots were probed on the same membrane with anti-mNeonGreen and anti-tubulin (loading control) antibodies. An additional western blot was performed with equal sample volumes and probed with anti-ANTXR2 antibody.

Given the high sequence identity of human and murine ANTXR2, we were intrigued that mANTXR2 did not constitute an efficient receptor. Since mANTXR2 differs the most from other mammalian orthologs in its Ig-like domain (Sup. Fig. 7), we generated chimeric proteins between hANTXR2 and mANTXR2, expressed the constructs in HAP1 cells and confirmed their correct cellular localization by IF (Fig. 4D, Sup. Fig. 6BC). Replacing the Ig-like domain of the human protein with the corresponding murine domain significantly decreased susceptibility of HAP1 cells to NetF (Figure 4EF). Conversely, replacing the murine Ig-like domain with the human Ig-like domain in mANTXR2 rendered cells susceptible to NetF. This suggests that, unlike for PA binding, the Ig-like domain of ANTXR2 is involved in NetF interaction and that differences between the murine and human protein in this domain affect this interaction.

### Structural characterization of NetF and ANTXR2 interaction

We overexpressed the extracellular region of hANTXR2 (termed extANTXR2) in Expi293 cells, incorporating a C-terminal FLAG and Hisx6 tag (Sup. Fig. 8A). Addition of purified extANTXR2 to pre-formed NetF oligomers in detergent resulted in receptor binding with varying stoichiometries, most commonly two to four extANTXR2 molecules per NetF pore, as observed by cryo-EM (Sup. Fig. 8). Since binding on consecutive protomers was commonly observed without apparent steric hindrance, we hypothesize that, *in vivo*, the ANTXR2-NetF protomer stoichiometry within a NetF oligomer could reach up to a 1:1 ratio. Particle classification of the consensus structure (Figure 5A, Sup. Fig. 8E) revealed subsets with ANTXR2 molecules bound that could achieve sufficiently high resolution for detailed structural characterization of the interaction (Sup. Fig. 8F). Symmetry expansion and focused refinement on a single NetF protomer with bound ANTXR2 further improved the resolution of the receptor binding site (Figure 5C).

**Figure 5.**
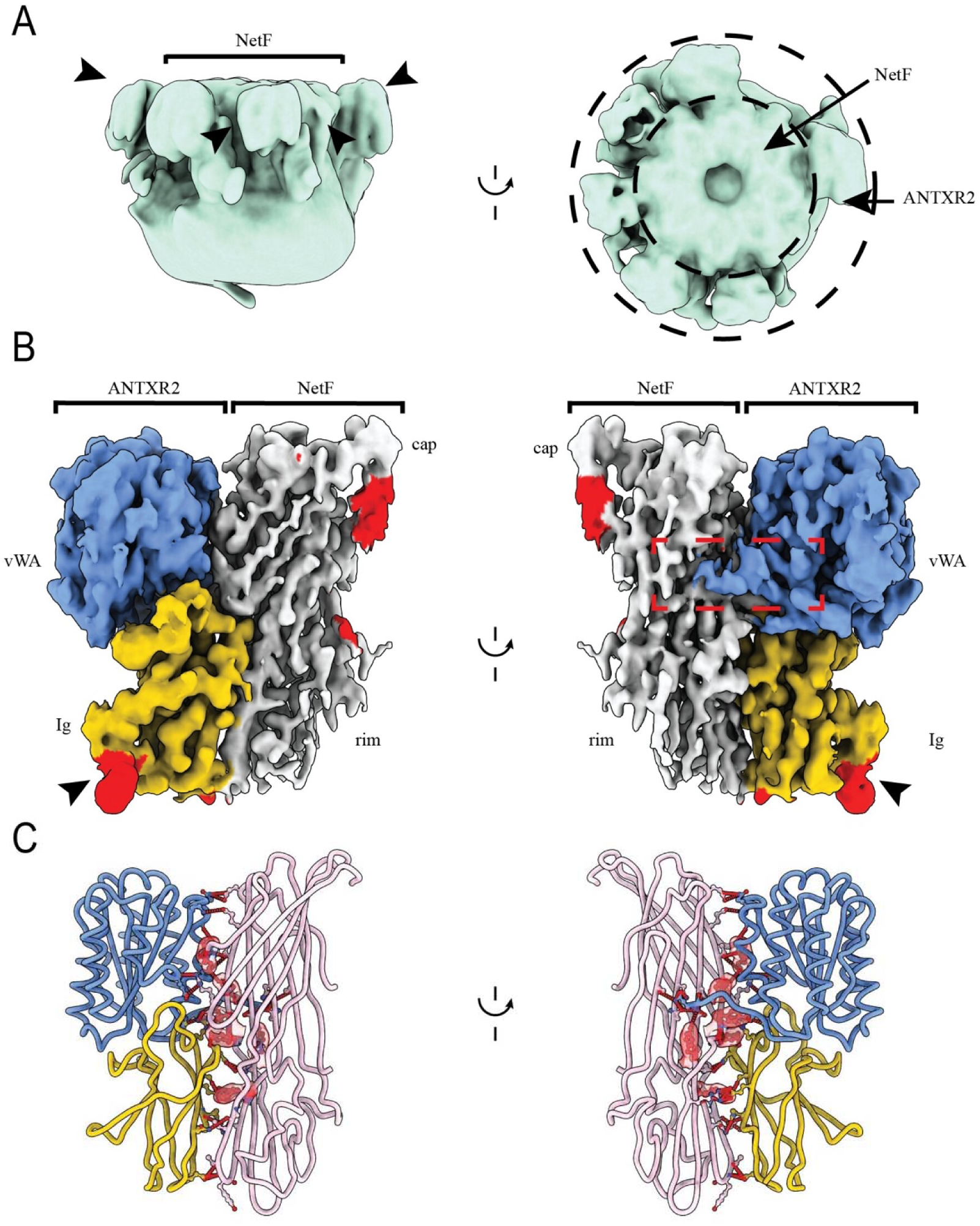
Structural characterization of the NetF ANTXR2 interaction. **(A)** Top and side views of the consensus initial model obtained from initial processing of NetF:ANTXR2 sample. The density of NetF is marked as the inner density while the position of ANTXR2 density is marked by black arrows. **(B)** Local resolution cryo-EM map of NetF protomer bound to ANTXR2. NetF density is shown in gray while for ANTXR2 the vWA domain is shown in blue and the Ig-like domain in yellow. Extra densities corresponding to the NetF N-terminus and the glycosylation of Asp260 (black arrow) are shown in red. Two representations rotated by 180° are shown for clarity. The red dotted rectangle highlights protein – protein contacts of the N-terminus of ANTXR2 with NetF. **(C)** The interface between NetF and ANTXR2 is shown in cartoon representation. Residues involved in the interface are shown as ball-and-stick representation and the distances are shown as red dashed lines. NetF is shown in pink while for ANTXR2 the vWA domain is shown in blue and the Ig-like domain is shown in yellow. The tyrosine patch at the interface is shown as transparent surface representation in red. Two views rotated by 180° are shown for clarity.

The 2.1 – 3.12 Å resolution map shows that NetF interacts with ANTXR2 over a contact area of approximately 1400 Å^2^, involving binding sites on both VWA and Ig-like domains (Figure 5C, Table S1 and S3). A single ANTXR2 molecule interacts with only a single NetF protomer. The contact site comprises more than 40 amino acids from each protomer, with hydrogen bonds and salt bridges stabilizing the interaction (Figures 5C, Table S1 and S3). Notably, the N-terminus of ANTXR2 wraps around NetF protomer on one side of the cap/rim domain interface, while Leu 274 and Asn 275, located on a loop on the opposite side, are inserted in a groove between the same domains of NetF (Figure 5BC). A short loop of ANTXR2 (Gly 243 – Asn 250) which contains one of the two putative glycosylation sites (Asn250) is flexible and not well resolved in the EM map. Its position is however too far from the NetF protomer to be directly involved in binding. The glycosylation state of Asn 250 cannot be confidently determined, while a clear density at the second putative glycosylation site, Asn 260, confirms the presence of a glycan on the side of ANTXR2 facing away from the NetF-binding interface (Figure 5C). Overall, these data indicate that the two glycosylation sites Asn 250 and Asn 260 are not directly involved in the interaction with NetF. One particularity of the NetF structure is the presence of 15 exposed tyrosine residues on its surface with 9 of them on the outer cap and rim surface and 6 of them located in the contact site (Figure 5C). Hydrogen bonds and cation – pi interactions are part of the contact site. Six more tyrosine residues are involved in membrane interaction. This distribution suggests a role in receptor binding, potential oligomer stabilization, and membrane anchoring, highlighting tyrosine-rich regions as key structural features of NetF.

## DISCUSSION

In this study, we elucidate the structure of NetF in its oligomeric pre-pore and its membrane-inserted pore state. We show that it strongly interacts with lipids in a way reminiscent to the binding of Hla and other βPFTs of the hemolysin sub-family ^17,19,20^. In addition, we identify ANTXR2, as a receptor for NetF. By determining the co-structure of the NetF oligomeric pore bound to the extracellular domain of ANTXR2 we unravel the detailed structural basis of the interaction a hemolysin βPFT and its membrane receptor.

Our results show that although anthrax PA and NetF share the same cellular receptor, their interaction with ANTXR2 markedly differs. PA binds ANTXR2 at the apical portion of the vWA domain, engaging the MIDAS motif, similarly to ANTXR2 interaction with collagen VI ^27,28^. In contrast, NetF interacts laterally with both the Ig-like and vWA domains of ANTXR2, making contacts across a broader surface area but without engaging with the MIDAS motif. Additionally, NetF interacts with membrane lipids through its rim domain, while also engaging with the membrane-proximal part of ANTXR2. During receptor binding, the toxin rim domain primarily interacts with the receptor Ig-like domain, while the cap domain interacts with the vWA domain. Another key difference that distinguishes NetF from anthrax PA is the length of their β-barrels. NetF forms a ∼50 Å-long β-barrel pore, matching the thickness of the lipid bilayer. In contrast, PA has an extended β-barrel (∼110 Å) because it binds at the distal region of ANTXR2, requiring it to span both the entire extracellular region of ANTXR2 (∼50 Å) and the lipid bilayer ^29^.

Prior to our study, the only reported structure of a hemolysin βPFT in complex with the extracellular portion of its receptor was that of *S. aureus* LukGH bound to CD11b/CD18 ^30^. Interestingly, LukGH interacts with the most distal part of CD11b, approximately 200-250 Å away from the membrane when the receptor is fully extended. Given that its β-barrel length is similar to that of NetF, LukGH pore formation would require either receptor bending or toxin detachment from the receptor. In contrast, NetF binds to the membrane-proximal Ig-like domain of ANTXR2, enabling direct membrane interaction and pore insertion. Interesting, like NetF, CPB binds the membrane-proximal Ig-like domain of its receptor, CD31 ^21^, suggesting a conserved binding pattern among clostridial hemolysin βPFTs. Additionally, the ANTXR2 structure obtained in this study is the first structure of ANTXR2 complete extracellular domain, including its posttranslational modifications.

While we elucidated the binding mechanism of NetF, its role in disease development remains to be determined. We show that NetF effectively targets ANTXR2 of various mammals, including dogs and horses, where NetF-producing *C. perfringens* strains are linked to enteric disease ^10^. Since anthrax toxin induces immune suppression and vascular collapse ^31–34^, primarily by targeting ANTXR2 ^35^, NetF may play a similar role in canine and equine enteritis. However, unlike anthrax PA, NetF does not use ANTXR1 as a receptor. In addition, our data suggests that while NetF is able to bind directly to lipids and oligomerize even in the absence of ANTXR2 it does not lead to toxicity and cell death. Moreover, while our data show different binding specificities towards different species – particularly murine – the residues identified in the protein-protein interaction appear to be mostly conserved.

In conclusion, by revealing the extensive molecular interaction of NetF with the vWA and Ig-like domains of its receptor ANTXR2, we enhance our understanding of receptor-mediated targeting by hemolysin βPFTs and provide unprecedented insight into the specific amino acid interactions driving toxin-receptor binding.

## MATERIALS AND METHODS

### RESOURCE AVAILABILITY

#### Lead Contact

Further information and requests for resources and reagents should be directed to and will be fulfilled by the lead contact, Benoît Zuber.

#### Materials Availability

This study did not generate new unique reagents. Plasmids generated in this study are available from the corresponding authors upon request.

#### Data and Code Availability

RNA-seq data have been deposited at the European Nucleotide Archive (ENA, accession number: *PRJEB78819*) and are publicly available as of the date of publication.

Cryo-EM maps and their respective models are available in the EMDB: NetF prepore in MSP2N2 nanodiscs: PDB: 9RSM ; EMDB: EMD-54221 NetF pore in DDM:CHS: PDB: 9RSU; EMDB: EMD-54226

NetF-ANTXR2: PDB: 9RT2, 9RT4; EMDB: EMD-54238, EMD-54245

### EXPERIMENTAL MODEL

#### Cell Culture

HEK293FT, HFF-1, EpH4, bEnd.3, equine and canine primary fibroblasts, J3T-bg, MDCK, and BoMac cell lines were cultured in DMEM medium (Gibco, #41965039) supplemented with 10% fetal calf serum FCS (BioConcept, #2-01F00-I) and 2 mM L-Glutamine (Gibco). HeLa and ECaNep cell lines were cultured in MEM medium (Gibco, #21090022), supplemented with 10% FCS and 2 mM L-Glutamine. U937 cells were cultured in RPMI 1640 medium (Gibco, #11875093) supplemented with 10% FCS and 2 mM L-Glutamine. HAP1 cells were cultured in IMDM medium with GlutaMAX™ (Gibco, #31980022) supplemented with 10% FCS. BUcEC cells were cultured in PriGrow C medium (Abm, #TM100) supplemented with 2% FCS. All cell lines were cultured in the presence of penicillin-streptomycin (Gibco) and grown at 37°C in an atmosphere containing 5% CO2.

### METHOD DETAILS

#### Plasmids

All plasmids used in this study are listed in Table S4. psPAX2 was a gift from Didier Trono (Addgene plasmid # 12260 ; http://n2t.net/addgene:12260 ; RRID:Addgene_12260). pMD2.G was a gift from Didier Trono (Addgene plasmid # 12259 ; http://n2t.net/addgene:12259 ; RRID:Addgene_12259). LentiCRISPR v2 was a gift from Feng Zhang (Addgene plasmid # 52961 ; http://n2t.net/addgene:52961 ; RRID:Addgene_52961) ^36^. pHR-SFFV_3C-Twin-Strep was a gift from A. Radu Aricescu (Addgene plasmid # 113900 ; http://n2t.net/addgene:113900 ; RRID:Addgene_113900) ^37^. pMSP2N2 was a gift from Stephen Sligar (Addgene plasmid # 29520 ; http://n2t.net/addgene:29520 ; RRID:Addgene_29520) ^38^.

#### Primers and sgRNAs

All primers and sgRNAs used in this study were ordered from Microsynth AG (Balgach, Switzerland) and are listed in Table S5.

#### Recombinant Protein Expression and Purification

Hexahistidine tagged NetF in pET-19b was expressed in E. Coli BL21(DE3).50-100 ng plasmid was transformed in 50 µl bacteria by heat shock (45 seconds at 42°C) and plated on selection plates with 100 µg/ml ampicillin. A single colony was used to inoculate a starting pre-culture of 100 ml LB with ampicillin. The pre-culture was allowed to grow overnight at 37°C. Cultures for protein expression were started by adding 5 ml of the pre-culture into 500 ml LB with ampicillin and grown at 37°C until OD600 of 0.6-0.8, when temperature was decreased to 18°C. Once the culture reached 18°C protein expression was induced by the addition of 100 µM IPTG. Expression was carried out overnight, followed by centrifugation. Bacterial pellets were resuspended in 50 mM Tris pH 8.0, 500 mM NaCl, 20 mM Imidazole, 2 mM β-mercaptoethanol supplemented with EDTA-free protease inhibitors (Sigma), lysozyme, 5 mM MgCl_2_ and benzonase. Cells were broken by three passages through an LM10 microfluidizer (Microfluidics) and the cell lysate was cleared by centrifugation at 25,000 x g for one hour at 4 °C. The supernatant was loaded on a 5 ml HisTrap column (Cytiva) and eluted using a 0 to 500 mM imidazole gradient. Fractions containing NetF were pooled and dialyzed against 20 mM Tris pH 8, 150 mM NaCl, 1 mM Dithiothreitol. To obtain pure NetF monomers, the sample was run on a Superdex 200 Increase 10/300 GL (Cytiva) size exclusion chromatography column. The purity of the sample was checked by SDS-PAGE.The extracellular domain of ANTXR2 (amino acids 36-282) was cloned for expression in Expi293 cells (Thermo Fischer) with a N-terminal secretion signal and a C-terminal hexahistidine tag followed by a FLAG tag. The vector was a kind gift of Sylvia Ho. For transfection Expi293 cells were grown in Expi293 expression medium (Gibco). Cells were seeded at 2×10^6^ cells/ml in fresh medium and allowed to grow overnight at 37°C, 8% CO_2_. 126 µg of plasmid was added to 5 ml Opti-MEM medium (Gibco) and vortexed. Polyethylnimine (Polysciences) was added at final concentration of 75 µg/ml. The mix was incubated at room temperature for 15-20 minutes and added to the cells. Cells were placed in the incubator and aliquots were removed daily to check protein expression levels and viability. After 72 hours, soy hydrolysate 1.6 ml (50x stock), glucose 0.54 ml (45% stock) and fresh media (17.86ml) were added as supplements and the cells were grown for another four days before harvesting. The supernatant from 60 ml culture was collected and incubated with 1 ml of anti-FLAG M2 magnetic beads (Sigma) at 4°C overnight. Subsequently the beads were washed three times with TBS (50 mM Tris pH 7.5, 50 mM NaCl) followed by three elutions with 1 ml TBS with 100 µg/ml FLAG peptide (Sigma). The different fractions were inspected by SDS-PAGE and the fractions containing ANTXR2 were pulled and concentrated to 0.3 mg/ml.

#### Preparation of NetF oligomers

Liposomes and MSP2N2 nanodiscs were prepared as previously described ^6,39^. Briefly the respective lipids were mixed in chloroform and dried under nitrogen followed by one hour under high vacuum. The lipid film was solubilized in 50mM Tris pH 7.5, 500 mM NaCl for liposomes and 20 mM Hepes pH 8, 100 mM KCl, 100 mM Na cholate for nanodiscs. Liposomes were extruded through a 100 nm filter (Avanti) followed by incubation with NetF (1:166 molar ratio). Oligomerization on liposomes was checked by SDS-PAGE and the oligomers were solubilized in DDM:CHS 1:0.1%. MSP2N2 was expressed in *E. coli* and purified on HisTrap column (Cytiva) as described above. Nanodiscs were formed by detergent removal with biobeads (biorad) followed by dialysis against 20 mM Hepes pH 7.5, 100 mM KCl overnight. NetF was added directly to preformed nanodiscs and checked by cryo-EM (1:10 molar ratio).

#### Sample preparation for cryo-EM

All samples for cryo-EM were vitrified using an FEI Vitrobot 4 according to the manufacturer’s instructions. The grids (Quantifoil 2/1 or 1.2/1.3 with 2 nm C) were glow discharged prior to usage. After vitrification, the grids were screened on an FEI Tecnai F20 equipped with a Falcon III detector and Gatan 626 cryo-holder. For high resolution acquisition, the grids were clipped and loaded directly into a FEI Titan Krios G4 equipped with a Falcon 4i detector and Selectris energy filter with a 20 eV slit.

#### Single particle acquisition, reconstruction, and atomic model building

Data was acquired at 165000x magnification corresponding to a pixel size of 0.73 Å^2^. Automatic data acquisition was set up using EPU automatic mode as detailed in Sup. Fig. 9 and Sup. Fig. 10 and saved as eer stacks with a total dose of ∼40e^-^/Å^2^. The movies were converted to tiff stacks using relion ^40^ and processed in cryosparc ^41^. Briefly, motion correction ^42^ and patch CTF^43^ was performed followed by template picker. The automatic picks were visually inspected, the particles were extracted and 2D classified. Following several rounds of 2D classification a subset of the particles was used to generate an initial model followed by homogeneous refinement. In the case of the ANTXR2 complex with NetF, after a 3D refinement procedure, the dataset was subjected to 3D classification, and all particles belonging to classes exhibiting receptor density were subsequently pooled (Sup. Fig. 8).. Symmetry expansion was performed followed by a new round of 3D classification with a focused mask around one NetF protomer and ANTXR2 density. The signal was subtracted from the dataset and local refinement was performed to obtain the map of ANTXR2 bound to a NetF protomer. ModelAngelo ^44^ was used for automated model building followed by Phenix ^45^ dock, rebuild and real-space refine and manual inspection and corrections in C*oot* ^46^. To calculate the pore diameter and generate the figure 2D we used HOLE ^47^ and hole-cmm ^48^ to obtain a ChimeraX-compatible model.

#### Cytotoxicity assays

NetF-induced cytotoxicity was evaluated using a Resazurin-based viability assay. 50,000 U937 cells in suspension were plated per well of a 96 well plate. Fibroblasts (HFF-1, equine and canine primary fibroblasts) were grown to 100% confluency in a 96 well plate. All other cell lines were grown to 50% confluency in a 96 well plate. All cells were then incubated with NetF for 24h. Resazurin salt (Sigma, #R7017) was diluted in PBS and added to cells at a 0.002% final concentration, incubated for 2h at 37°C and fluorescent signal intensity was quantified using an Hidex Sense microplate reader (Hidex Oy; 544/590nm Ex/Em). Signal intensity of NetF-treated cells was normalized to that of untreated control cells to determine relative viability.

#### RNA-seq

NetF-resistant U937 and J3T subclones (U937-R, J3T-R) were obtained by keeping WT cells under continuous NetF pressure. U937 and J3T cell lines were cloned by limiting dilution to obtain single-cell-derived populations. Single-cell clonal populations were treated with 1 µg/ml NetF for 24h. Surviving cells were then cultured in medium freshly supplemented with 1 µg/ml NetF for the following 14 days to obtain complete NetF-resistant cell populations. Total RNA of WT and NetF-resistant cells was isolated using a RNeasy Mini Kit (Qiagen, #74104) following the manufacturer’s instruction. RNA purity was assessed by spectrophotometry (NanoDrop) and RNA integrity was assessed with a 2100 BioAnalyzer (Agilent). PolyA selected mRNAs were sequenced at Novogene Cambridge Sequencing Centre. Differential expression analysis was performed using the edgeR R package (3.22.5). P values were adjusted using the Benjamini & Hochberg method. Corrected P-value of 0.05 and absolute fold change of 2 were set as the threshold for significantly differential expression.

#### Generation of ANTXR2 knockout cell lines

U937 and HFF-1 cells were transduced with lentiviral vectors expressing Cas9 and sgRNAs targeting ANTXR2 (sgRNA1 and sgRNA2; Table S4). sgRNAs were cloned into lentiCRISPR v2 and transformed into *E. coli* Endura cells (LGC Biosearch Technologies). Cells were grown at 30°C and plasmids were purified and Sanger sequenced with the U6 forward primer (5’-GGGCAGGAAGAGGGCCTAT-3’). Lentivirus was produced in HEK293FT cells by co-transfection of 22.5 μg transfer vector, 20 μg psPAX2, and 2 μg pMD2.G using calcium phosphate precipitation. After 5 min incubation with 50 μL of 2.5LM CaCl₂, the DNA was mixed with 2× HEPES-buffered saline and added dropwise to 2 × 10L HEK293FT cells plated the previous day in 100 mm dishes. Growth medium was replaced 16 h post-transfection. Lentivirus-containing medium was collected at 40 h and 64 h post-transfection, filtered through 0.45Lμm cellulose filters, and stored at –80 °C. For transduction, cells were seeded in T25 flask at a density yielding ∼20% confluency in 4 ml viral supernatent with 8 μg/ml Polybrene (Sigma) and incubated for 24h Puromycin selection was initiated 24 h post-transduction (2 μg/ml for U937, 1 μg/ml for HFF-1) and maintained for 3 days. Knockout efficiency was assessed using the TIDE web tool ^49^. Genomic DNA of each cell line was extracted using the DNeasy Blood & Tissue Kit (Qiagen, #69504), and target loci were PCR-amplified using primers flanking the sgRNA sites. Amplicons were purified (NucleoSpin Gel and PCR Clean-up, Macherey-Nagel) and Sanger sequenced (Microsynth, Switzerland)..

#### Generation of transgenic HAP1 and HEK293 cells

C-terminally mNeonGreen-tagged constructs were generated by cloning ANTXR2 and TEM8 variants (Table S5) into a lentiviral transfer vector under the control of a CMV promoter, with a puromycin resistance cassette for selection. Murine, canine, equine, and bovine ANTXR2 sequences, as well as chimeric and truncated constructs, were derived from human and murine templates using standard cloning procedures. Site-directed mutagenesis was used to introduce point mutations and small region deletions. Primers used for all constructs are listed in Table S5. All plasmids were sequence-verified. Lentivirus was produced in HEK293FT cells using a second-generation packaging system and calcium phosphate transfection. Viral supernatants were collected at 40 and 64 h post-transfection, filtered, and used to transduce HAP1 and HEK293 cells. Puromycin selection (0.8 μg/ml for HAP1, 1.5 μg/ml for HEK293) was applied for 3 days. Due to low basal expression in HAP1 cells, transduced populations were FACS-sorted to isolate the top 5% mNeonGreen-expressing cells. Monoclonal HAP1 populations expressing selected constructs (Table S6) were further isolated by limiting dilution.

#### Immunoblotting

Cells were lysed in RIPA buffer (10 mM Tris-HCl pH 7.5, 150 mM NaCl, 0.5 mM EDTA, 0.1% SDS, 1 % Triton X-100, 1% Na-Deoxycholate) supplemented with protease inhibitor cocktail, and incubated for 30 min on ice. Lysates were clarified by centrifugation (14,000 x g, 15 min, 4 °C), mixed with reducing sample buffer to 2% SDS final concentration, and boiled for 5 min at 95°C. Proteins were resolved by SDS-PAGE and transferred on 0.45 mm nitrocellulose membranes (Thermo Scientific) using a Trans-Blot Turbo system (Bio-Rad). Membranes were blocked in 2% BSA for 1 h at room temperature and incubated overnight at 4°C with primary antibodies: rabbit anti-PA (1:2,000), mouse anti-β-tubulin (1:5,000), mouse anti-His-tag (1:1,000), rabbit anti-mNeonGreen (1:2,000), and mouse anti-ANTXR2 (1:2,000). Western blots with HRP were developed with Pierce™ ECL system (Thermo Scientific, #32209) and blots were scanned using Azure Biosystem gel imager.

#### Co-IP assays

Confluent cells were grown to confluency. For PA binding assays, lysates were diluted in 400 µl dilution buffer (10 mM Tris-HCl pH 7.5, 150 mM NaCl, 0.5 mM EDTA, protease inhibitors). Cells were incubated on ice with 1 µg/ml PA83 and 0.1 µg/ml lethal factor for 30 min. Afterwards, cells were lysed immediately or after 60 min incubation at 37°C to promote PA83 cleavage into PA63. For ANTXR2-Net-F binding assays, cells were incubated with 10 µg/ml NetF for 30 min on ice, followed by 15 min incubation at 37°C. All cells, including those detached due to NetF-induced toxicity, were collected for lysis.

Cells were washed with ice-cold PBS and lysed in RIPA buffer on ice for 30 min. Lysates were diluted with 400 μl dilution buffer (10 mM Tris-HCl pHL7.5, 150 mM NaCl, 0.5 mM EDTA, protease inhibitors) to reduce detergent concentrations. Clarified lysates (14,000 × g, 15Lmin, 4L°C for PA binding assays; 5,000 x g, 10 min, 4 °C for ANTXR2-Net-F binding assays) were either used directly for immunoblotting or incubated overnight with anti-mNeonGreen magnetic beads (Chromotek) according to the manufacturer’s instructions. Beads were washed three times with ice-cold buffer (10 mM Tris-HCl pHL7.5, 150 mM NaCl, 0.05% NP-40 substitute, 0.5LmM EDTA), resuspended in reducing sample buffer adjusted to 2% SDS, and boiled for 5Lmin at 95 °C. Beads were removed before performing immunoblotting as described above.

#### Fluorescence microscopy

Cells were seeded in 96-well imaging microplates (Agilent, #204626-100), grown to ∼50% confluency and fixed in 4% paraformaldehyde in PBS for 10 min at room temperature. After a PBS wash, cells were permeabilized in 0.2% Triton X-100 in PBS for 10 min, then incubated with 100 nM rhodamine-phalloidin (Cytoskeleton, #PHDR1) for 30 min. Following three PBS washes, DNA was stained with 0.1 µg/mL DAPI (Invitrogen) for 5 min. Cells were rinsed once in PBS and stored in PBS at 4°C until imaging. Images were acquired using a Nikon Ti2 Cicero spinning disc confocal microscope (40X objective). Further image analysis was performed using Fiji ^50^.

### QUANTIFICATION AND STATISTICAL ANALYSIS

Statistics were performed using GraphPad Prism 6. Unless otherwise specified, one-way ANOVA followed by Šidák’s multiple comparisons test was performed to assess pairwise group differences, with the family-wise error rate controlled at 5% (α = 0.05).

## Supporting information

All supplementary data

## ACKNOWLEDGEMENTS

We thank Prof. P. Plattet, University of Bern, for providing canine cell cultures. EM data were acquired on an instrument of the Dubochet Center for Imaging in Bern and supported by the Microscopy Imaging Center (MIC) of the University of Bern. We gratefully acknowledge Marek Kaminek and David Kalbermatter for their assistance with EM. We thank Sylvia Ho for providing the plasmids for protein expression in Expi293. This study was funded a University of Bern ID Grant (H.P., B.Z.), SNSF grant 310030_212837 (H.P.), and SNSF sinergia grant 10000175 (B.Z., H.P.).

## AUTHOR CONTRIBUTIONS

H.P. and B.Z. conceptualized the study, supervised the research, and secured funding.

C.W., F.C., I.I., and A.N. designed and performed experiments and analyzed the data.

F.F. and J.F. performed experiments.

L.A. performed experiments under the supervision of F.G.v.d.G.

F.C., I.I., and H.P. wrote the initial draft with input from C.W.

F.C., I.I., H.P., and B.Z. substantially revised and refined the manuscript.

All authors reviewed and approved the final version.

## Notes

### Competing Interest Statement

The authors have declared no competing interest.

## REFERENCES

1 Songer, J. G. in Handbook on Clostridia (ed J Dürre) Ch. 22, 527–544 (Taylor & Francis Group, 2005).

2 Songer, J. G. Clostridia as agents of zoonotic disease. Vet Microbiol 140, 399–404 (2010).

3 Uzal, F. A. et al. Comparative pathogenesis of enteric clostridial infections in humans and animals. Anaerobe 53, 11–20 (2018). 10.1016/j.anaerobe.2018.06.002

4 Popoff, M. R. & Bouvet, P. Clostridial toxins. Future Microbiol 4, 1021–1064 (2009).

5 Rood, J. I. et al. Expansion of the Clostridium perfringens toxin-based typing scheme. Anaerobe 53, 5–10 (2018).

6 Mehdizadeh Gohari, I., et al. A novel pore-forming toxin in type A Clostridium perfringens is associated with both fatal canine hemorrhagic gastroenteritis and fatal foal necrotizing enterocolitis. PLoS One 10, e0122684 (2015). 10.1371/journal.pone.0122684

7 Finley, A. et al. Prevalence of netF-positive Clostridium perfringens in foals in southwestern Ontario. Can J Vet Res 80, 242–244 (2016).

8 Leipig-Rudolph, M. et al. Intestinal lesions in dogs with acute hemorrhagic diarrhea syndrome associated with netF-positive Clostridium perfringens type A. J Vet Diagn Invest 30, 495–503 (2018). 10.1177/1040638718766983

9 Mehdizadeh Gohari, I., et al. NetF-positive Clostridium perfringens in neonatal foal necrotising enteritis in Kentucky. Vet Rec 178, 216 (2016). 10.1136/vr.103606

10 Mehdizadeh Gohari, I., Unterer, S., Whitehead, A. E. & Prescott, J. F. NetF-producing Clostridium perfringens and its associated diseases in dogs and foals. J Vet Diagn Invest 32, 230–238 (2020). 10.1177/1040638720904714

11 Sindern, N. et al. Prevalence of Clostridium perfringens netE and netF toxin genes in the feces of dogs with acute hemorrhagic diarrhea syndrome. J Vet Intern Med 33, 100–105 (2019). 10.1111/jvim.15361

12 Mehdizadeh Gohari, I., Brefo-Mensah, E. K., Palmer, M., Boerlin, P. & Prescott, J. F. Sialic acid facilitates binding and cytotoxic activity of the pore-forming Clostridium perfringens NetF toxin to host cells. PLoS One 13, e0206815 (2018). 10.1371/journal.pone.0206815

13 Dal Peraro, M. & van der Goot, F. G. Pore-forming toxins: ancient, but never really out of fashion. Nat Rev Microbiol 14, 77–92 (2016). 10.1038/nrmicro.2015.3

14 Jolivet-Reynaud, C., Hauttecoeur, B. & Alouf, J. E. Interaction of Clostridium perfringens delta toxin with erythrocyte and liposome membranes and relation with the specific binding to the ganglioside GM2. Toxicon 27, 1113–1126 (1989).

15 Bruggisser, J. et al. CD31 (PECAM-1) Serves as the Endothelial Cell-Specific Receptor of Clostridium perfringens beta-Toxin. Cell Host Microbe (2020). 10.1016/j.chom.2020.05.003

16 Huyet, J. et al. Structural Insights into Clostridium perfringens Delta Toxin Pore Formation. PLoS One 8, e66673 (2013). 10.1371/journal.pone.0066673

17 Savva, C. G. et al. Molecular architecture and functional analysis of NetB, a pore-forming toxin from Clostridium perfringens. J Biol Chem 288, 3512–3522 (2013). 10.1074/jbc.M112.430223

18 Abramson, J. et al. Accurate structure prediction of biomolecular interactions with AlphaFold 3. Nature, 1–3 (2024).

19 Galdiero, S. & Gouaux, E. High resolution crystallographic studies of alpha-hemolysin-phospholipid complexes define heptamer-lipid head group interactions: implication for understanding protein-lipid interactions. Protein Sci 13, 1503–1511 (2004). 10.1110/ps.03561104

20 Potrich, C. et al. The influence of membrane lipids in Staphylococcus aureus gamma-hemolysins pore formation. J Membr Biol 227, 13–24 (2009). 10.1007/s00232-008-9140-6

21 Bruggisser, J. et al. Cryo-EM structure of the octameric pore of Clostridium perfringens beta-toxin. EMBO Rep 23, e54856 (2022). 10.15252/embr.202254856

22 Choate, L. A. et al. Multiple stages of evolutionary change in anthrax toxin receptor expression in humans. Nat Commun 12, 6590 (2021). 10.1038/s41467-021-26854-z

23 Fedotova, A. A., Bonchuk, A. N., Mogila, V. A. & Georgiev, P. G. C2H2 Zinc Finger Proteins: The Largest but Poorly Explored Family of Higher Eukaryotic Transcription Factors. Acta Naturae 9, 47–58 (2017).

24 Gosztyla, M. L. et al. Integrated multi-omics analysis of zinc-finger proteins uncovers roles in RNA regulation. Mol Cell 84, 3826–3842 e3828 (2024). 10.1016/j.molcel.2024.08.010

25 Scobie, H. M., Rainey, G. J., Bradley, K. A. & Young, J. A. Human capillary morphogenesis protein 2 functions as an anthrax toxin receptor. Proc Natl Acad Sci U S A 100, 5170–5174 (2003). 10.1073/pnas.0431098100

26 Deuquet, J., Lausch, E., Superti-Furga, A. & van der Goot, F. G. The dark sides of capillary morphogenesis gene 2. EMBO J 31, 3–13 (2012). 10.1038/emboj.2011.442

27 Burgi, J. et al. CMG2/ANTXR2 regulates extracellular collagen VI which accumulates in hyaline fibromatosis syndrome. Nat Commun 8, 15861 (2017). 10.1038/ncomms15861

28 Lacy, D. B., Wigelsworth, D. J., Melnyk, R. A., Harrison, S. C. & Collier, R. J. Structure of heptameric protective antigen bound to an anthrax toxin receptor: a role for receptor in pH-dependent pore formation. Proc Natl Acad Sci U S A 101, 13147–13151 (2004). 10.1073/pnas.0405405101

29 Jiang, J., Pentelute, B. L., Collier, R. J. & Zhou, Z. H. Atomic structure of anthrax protective antigen pore elucidates toxin translocation. Nature 521, 545–549 (2015). 10.1038/nature14247

30 Trstenjak, N. et al. Molecular mechanism of leukocidin GH-integrin CD11b/CD18 recognition and species specificity. Proc Natl Acad Sci U S A 117, 317–327 (2020). 10.1073/pnas.1913690116

31 Baldari, C. T., Tonello, F., Paccani, S. R. & Montecucco, C. Anthrax toxins: A paradigm of bacterial immune suppression. Trends Immunol 27, 434–440 (2006). 10.1016/j.it.2006.07.002

32 Moayeri, M., Haines, D., Young, H. A. & Leppla, S. H. Bacillus anthracis lethal toxin induces TNF-alpha-independent hypoxia-mediated toxicity in mice. J Clin Invest 112, 670–682 (2003). 10.1172/JCI17991

33 Moayeri, M. & Leppla, S. H. The roles of anthrax toxin in pathogenesis. Curr Opin Microbiol 7, 19–24 (2004). 10.1016/j.mib.2003.12.001

34 Agrawal, A. et al. Impairment of dendritic cells and adaptive immunity by anthrax lethal toxin. Nature 424, 329–334 (2003). 10.1038/nature01794

35 Liu, S. et al. Capillary morphogenesis protein-2 is the major receptor mediating lethality of anthrax toxin in vivo. Proc Natl Acad Sci U S A 106, 12424–12429 (2009). 10.1073/pnas.0905409106

36 Sanjana, N. E., Shalem, O. & Zhang, F. Improved vectors and genome-wide libraries for CRISPR screening. Nature methods 11, 783–784 (2014).

37 Elegheert, J. et al. Lentiviral transduction of mammalian cells for fast, scalable and high-level production of soluble and membrane proteins. Nat Protoc 13, 2991–3017 (2018). 10.1038/s41596-018-0075-9

38 Grinkova, Y. V., Denisov, I. G. & Sligar, S. G. Engineering extended membrane scaffold proteins for self-assembly of soluble nanoscale lipid bilayers. Protein Eng Des Sel 23, 843–848 (2010). 10.1093/protein/gzq060

39 Ritchie, T. K. et al. Chapter 11 - Reconstitution of membrane proteins in phospholipid bilayer nanodiscs. Methods Enzymol 464, 211–231 (2009). 10.1016/S0076-6879(09)64011-8

40 Scheres, S. H. A Bayesian view on cryo-EM structure determination. J Mol Biol 415, 406–418 (2012). 10.1016/j.jmb.2011.11.010

41 Punjani, A., Rubinstein, J. L., Fleet, D. J. & Brubaker, M. A. cryoSPARC: algorithms for rapid unsupervised cryo-EM structure determination. Nat Methods 14, 290–296 (2017). 10.1038/nmeth.4169

42 Zheng, S. Q. et al. MotionCor2: anisotropic correction of beam-induced motion for improved cryo-electron microscopy. Nat Methods 14, 331–332 (2017). 10.1038/nmeth.4193

43 Rohou, A. & Grigorieff, N. CTFFIND4: Fast and accurate defocus estimation from electron micrographs. J Struct Biol 192, 216–221 (2015). 10.1016/j.jsb.2015.08.008

44 Jamali, K. et al. Automated model building and protein identification in cryo-EM maps. Nature 628, 450–457 (2024). 10.1038/s41586-024-07215-4

45 Liebschner, D. et al. Macromolecular structure determination using X-rays, neutrons and electrons: recent developments in Phenix. Acta Crystallogr D Struct Biol 75, 861–877 (2019). 10.1107/S2059798319011471

46 Emsley, P., Lohkamp, B., Scott, W. G. & Cowtan, K. Features and development of Coot. Acta Crystallogr D Biol Crystallogr 66, 486–501 (2010). 10.1107/S0907444910007493

47 Smart, O. S., Neduvelil, J. G., Wang, X., Wallace, B. A. & Sansom, M. S. HOLE: a program for the analysis of the pore dimensions of ion channel structural models. J Mol Graph 14, 354–360, 376 (1996). 10.1016/s0263-7855(97)00009-x

48 joelmeyerson. hole-cmm, <https://github.com/joelmeyerson/hole-cmm> (2025).

49 Brinkman, E. K., Chen, T., Amendola, M. & Van Steensel, B. Easy quantitative assessment of genome editing by sequence trace decomposition. Nucleic acids research 42, e168–e168 (2014).

50 Schindelin, J., et al. Fiji: an open-source platform for biological-image analysis. Nat Methods 9, 676–682 (2012). 10.1038/nmeth.2019

