## Supplementary material for "Anthrax toxin receptor 2 is the Receptor for *Clostridium perfringens* NetF: Structural Insights into Toxin Binding and Pore Formation": All supplementary data

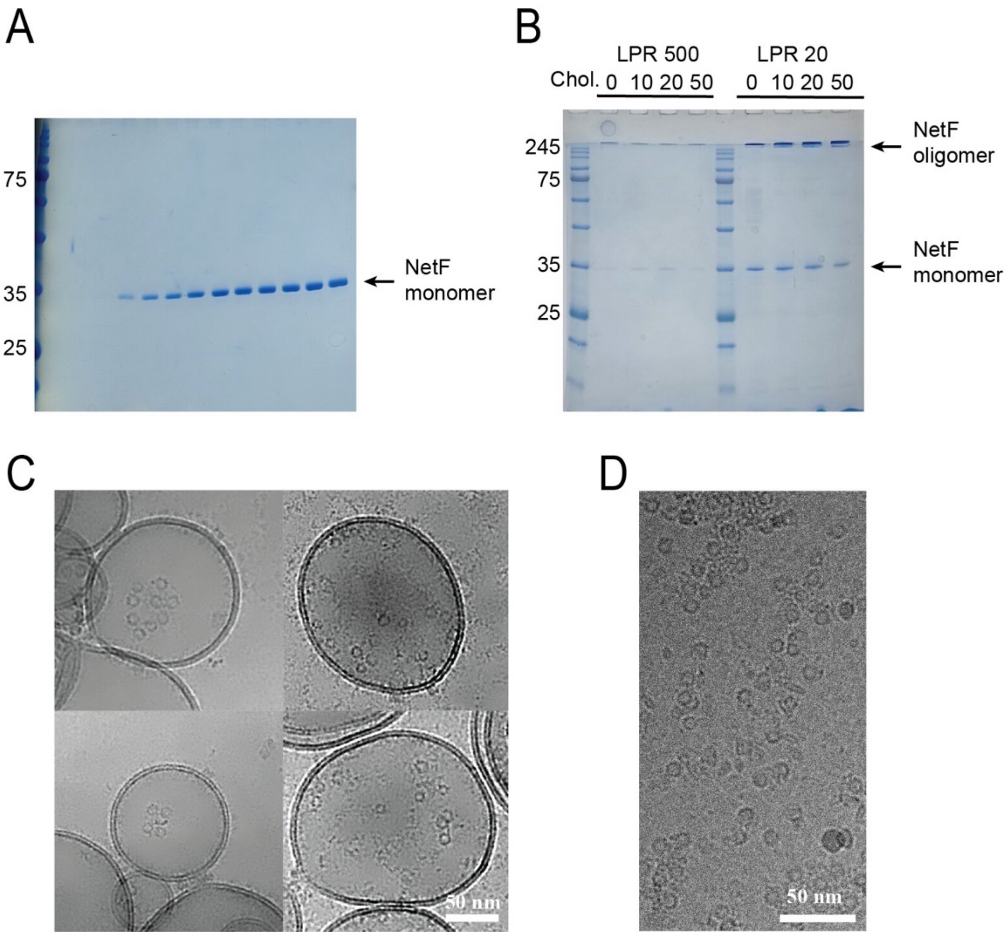


**Sup. Fig. 1.**

**(A)** Imidazole gradient elution fractions of the purification of NetF.

**(B)** Oligomerization of NetF at 500 (0.25µM NetF) and 20 (3µM NetF) lipid-to-protein ratio (LPR) on DOPC:DOPG liposomes containing the specified amount of cholesterol (0 to 50%).

**(C)** Cryo-EM images of NetF oligomers formed on DOPC:DOPG:Cholesterol liposomes.

**(D)** Cryo-EM image of NetF oligomers formed on DOPC:DOPG:Cholesterol MSP2N2 nanodiscs.

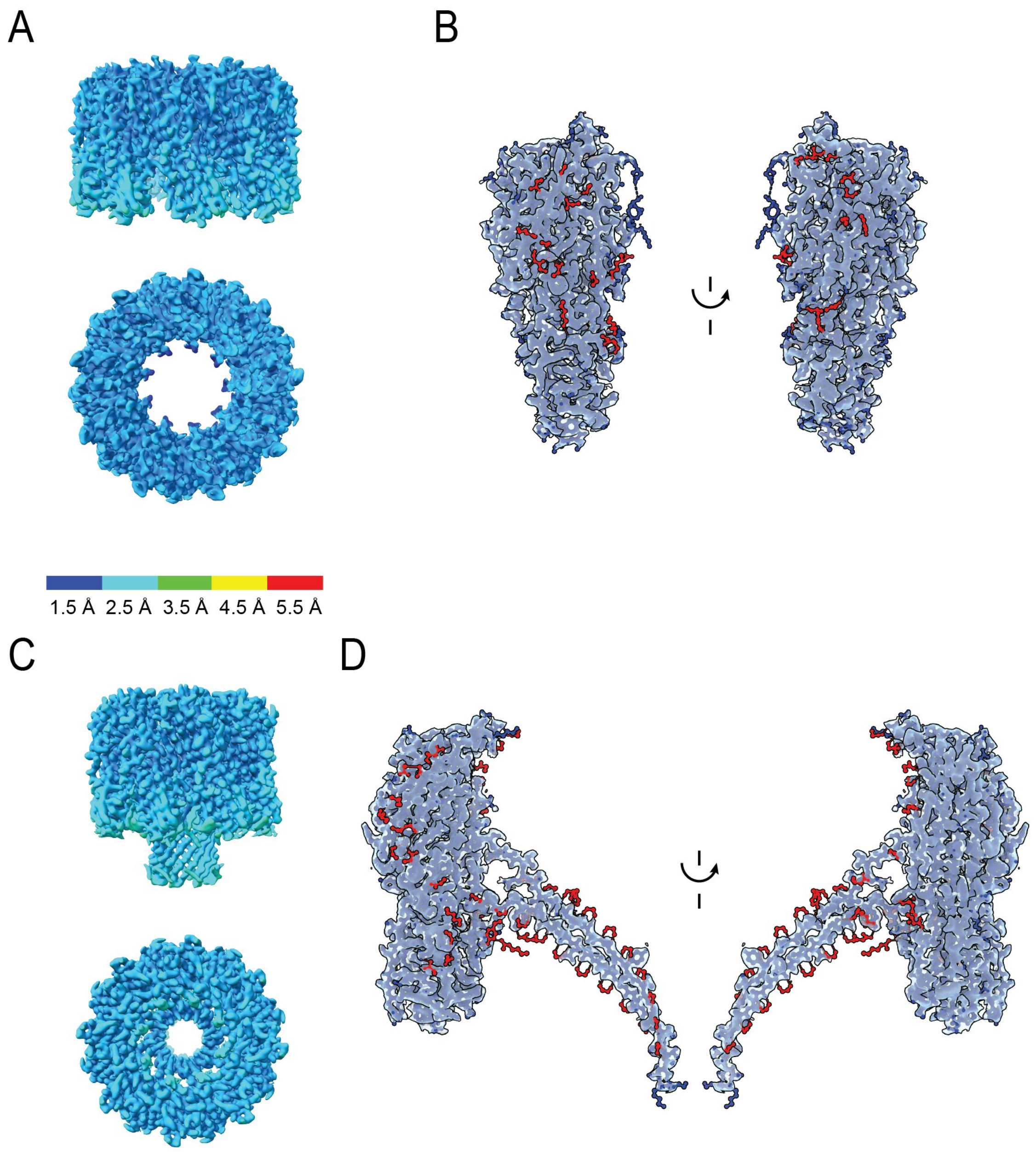


**Sup. Fig. 2.**

**(A)** Cryo-EM map of NetF in pre-pore conformation (C9 symmetry) colored coded by local resolution. Global resolution of the map 2.1 Å.

**(B)** Ball and stick representation of one NetF protomer (blue) docked in the cryo-EM map. In red residues from the neighboring protomers involved in the contact interface are shown while the dashed line shows the location of the unresolved residues in the pre-stem loop. Two representations rotated by 180° are shown for clarity.

**(C)** Cryo-EM map of NetF in pore conformation (C8 symmetry) colored coded by local resolution. Global resolution of the map 2.2 Å.

**(D)** Ball and stick representation of one NetF protomer (blue) docked in the cryo-EM map. In red residues from the neighboring protomers of the protomer – protomer interface are shown. Two representations rotated by 180° are represented for clarity.

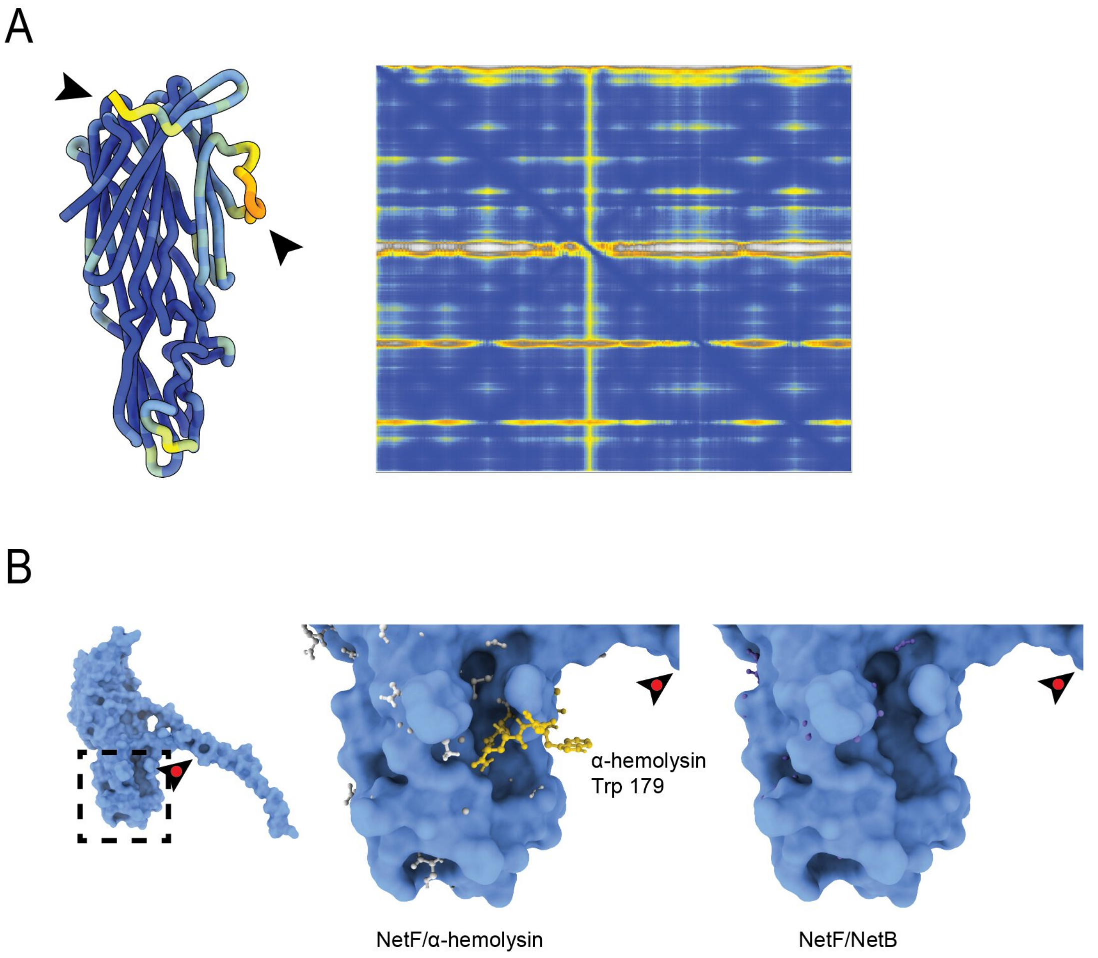


**Sup. Fig. 3.**

**(A)** Alphafold3 prediction of NetF monomer in its soluble conformation (left) colored by the pLDDT plot (right) indicating the per-residue local confident. The two domains with lower confidence, the N-terminus (left) and the pre-stem loop (right) are indicated by black arrows.

**(B)** Surface representation of one NetF protomer extracted from the pore model (left) highlighting the rim domain (dotted square). The position of the stem domain is shown by a black arrow with a red dot in all panels. View of the rim domain (middle) with a ball and stick representation of a *S. aureus* α-hemolysin protomer (gray) docked. The extra loop present in α-hemolysin containing tryptophane 179 is shown in yellow. In contrast, docking *C. perfringens* NetB docked in the NetF surface representation (right) shows no extra loops or residues in the rim domain.

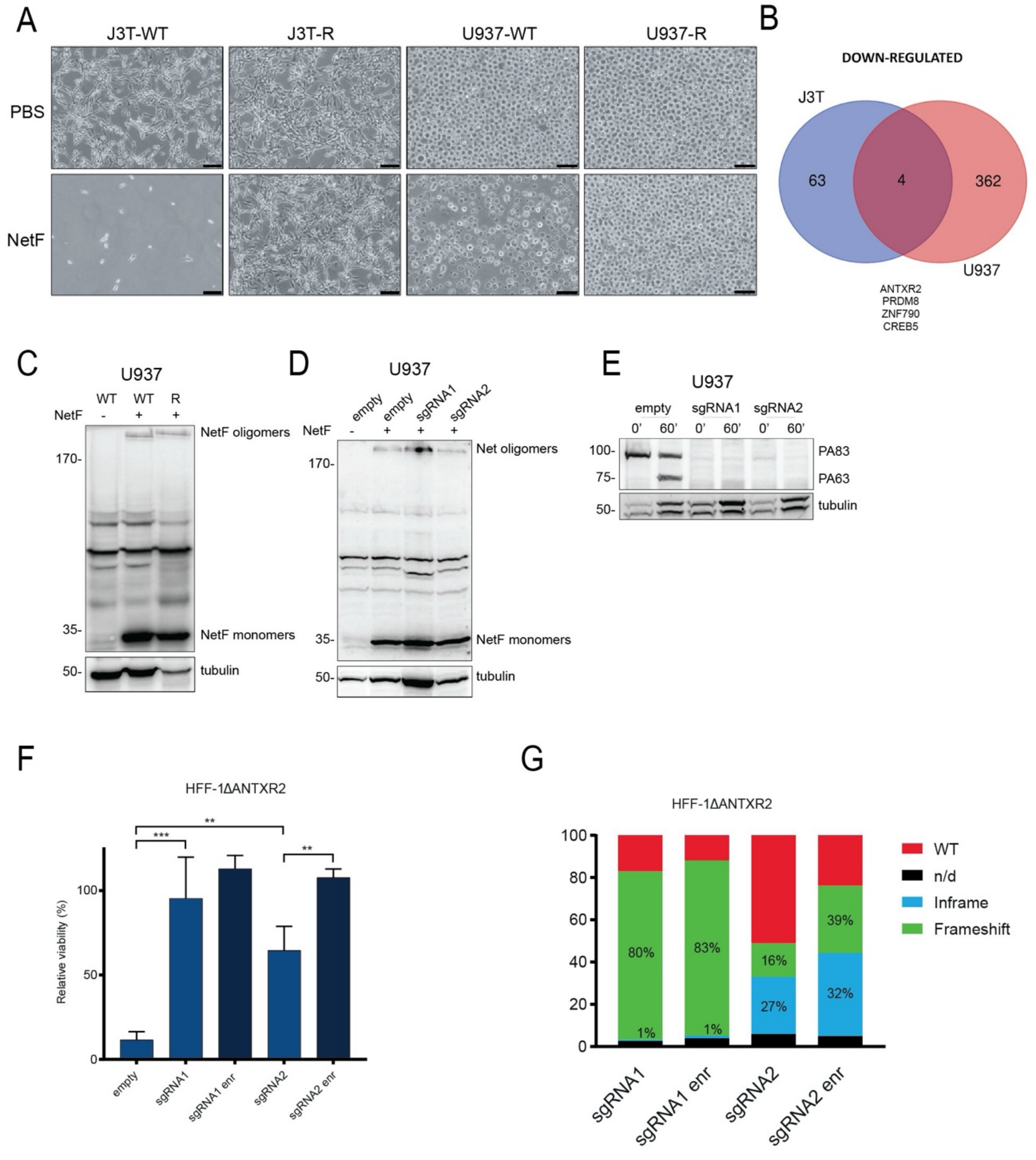


**Sup. Fig. 4.**

**(A)** Micrographs of J3T and U937 wild type (WT) and NetF-resistant (R) subpopulations treated with 1 µg/ml NetF (24h, 37°C).

**(B)** Venn's diagram summarizing the number of downregulated genes in U937 and J37 RNAseq experiments. ANTXR2 is the only gene encoding for a transmembrane protein.

**(C)** Western blot analysis of NetF oligomerization assay performed with U937-WT and U937-R cells. Cells not treated with NetF (-) were used as negative control. Immunoblot was performed on the same membrane using anti-His and anti-tubulin (loading control) antibodies. Figure shows one of three replicates of the same experiment.

**(D)** Western blot analysis of NetF oligomerization assay performed with U937 cells treated with sgRNA1 and sgRNA2, targeting the ANTXR2 gene, as well as an empty vector (empty). Cells not treated with NetF (-) were used as negative control. Experiment was performed as in S5A. Figure shows one of three replicates of the same experiment.

**(E)** Western blot analysis of PA binding assays performed with U937 cells treated with sgRNA1 and sgRNA2, targeting the ANTXR2 gene, as well as an empty vector (empty). Experiment was performed as in S5A. Figure shows one of three replicates of the same experiment.

**(F)** Viability of HFF-1 cells targeted by sgRNA1- and sgRNA2-mediated ANTXR2 knockout after incubation with 1 µg/ml NetF (24 h, 37°C), in % to untreated control cells. Mutations-enriched (enr.) populations were obtained by applying selective pressure with 1 µg/ml NetF, two times (2x 24 h, 37 °C) before proceeding with cytotoxicity assays. Data are represented as means (n=9) ± SD. One-way ANOVA, Sidak's multiple comparison test, *** ($p<0.001$), ** ($p < 0.01$).

**(G)** TIDE analysis showing high percentage of frameshift and in frame mutations in the ANTXR2 gene after sgRNA1- and sgRNA2-mediated ANTXR2 knockout in HFF-1 cells. Enrichment (enr.) of these mutations was obtained by applying two times selective pressure with 1 µg/ml NetF (2x 24 h, 37 °C) before proceeding with DNA extraction.

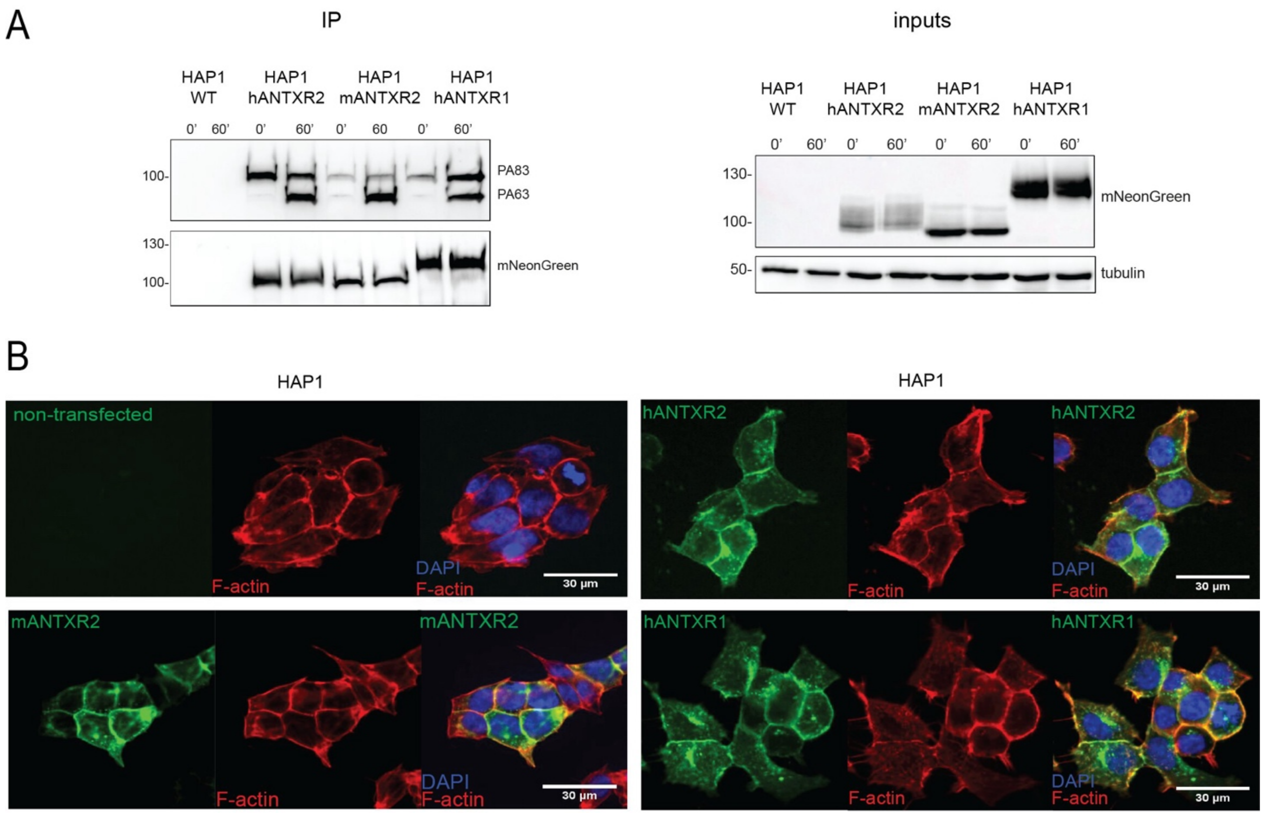


**Sup. Fig. 5.**

**(A)** Westen blot analysis of co-IP experiments in HAP1 cells expressing human ANTXR2, mouse ANTXR2 and human ANTXR1 incubated with 1 µg/ml PA on ice for 30 minutes and then either directly lysed (0') or incubated at 37°C for 1 hour (60'). IP was performed using anti-mNeonGreen beads and showed successful pull-down of PA with all three receptors (IP), demonstrating their functional expression at the plasma membrane. Immunoblot containing 50% eluate (IP) was probed on the same membrane with anti-PA antibody and anti-mNeonGreen antibody (pulldown efficiency control). Immunoblot containing 5% loading fraction (input) was probed on the same membrane using anti-mNeonGreen antibody and anti-tubulin antibody (loading control). Figure shows one of two replicates of the same experiment. Fluorescent microscopy pictures showing cellular localization of mNeonGreen-tagged hANTXR2, mANTXR2, and hANTXR1 in HAP1 cells. Rhodamine-phalloidin stained F-actin used as proxy for plasma membrane localization. Nuclei were stained with DAPI. Top left: untransfected cells as control.

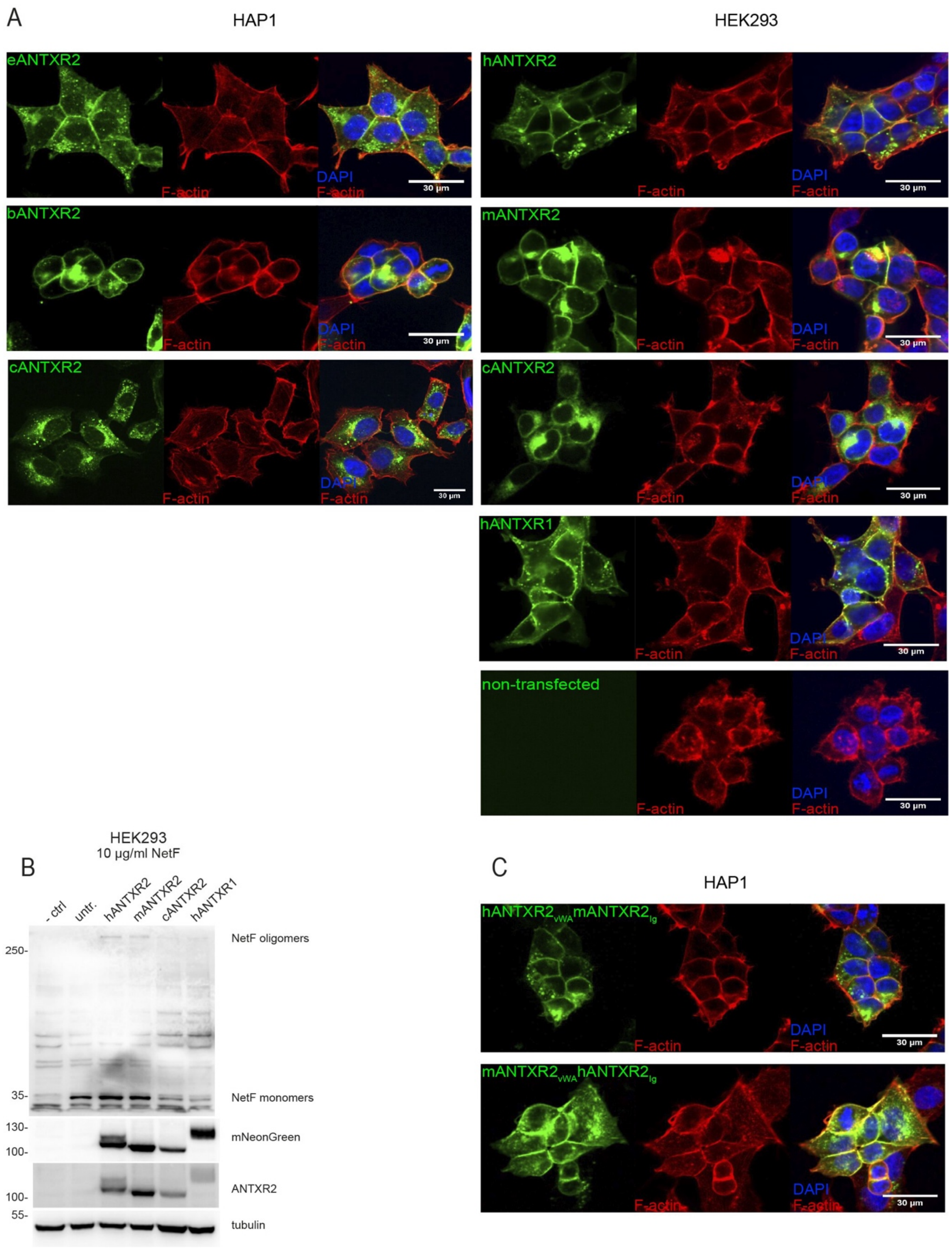


**Sup. Fig. 6.**

**(A)** Fluorescent microscopy pictures showing cellular localization of indicated mNeonGreen-tagged constructs in HAP1 and HEK293 cells. h: human, e: equine; b: bovine; c: canine, m: murine. Note that c ANTXR2 showed increased intracellular signal indication ER retention in both cell lines but also detectable membrane expression in HEK293. Staining performed as in Supl. Fig. 5.

**(B)** Western blot analysis of NetF oligomerization assay performed by incubating NetF at high concentrations (10 µg/ml) with HEK293 cells, untransduced (untr.) or overexpressing human, mouse and canine ANTXR2, and human ANTXR1. Immunoblot was performed on the same membrane using anti-His, anti-mNeonGreen and anti-tubulin (loading control) antibodies. An additional immunoblot was performed with equal sample volumes and probed with anti-ANTXR2 antibody.

**(C)** Fluorescent microscopy pictures showing cellular localization of mNeonGreen-tagged human and mouse ANTXR2 extracellular domain chimeras. Staining performed as in Fig. S5.

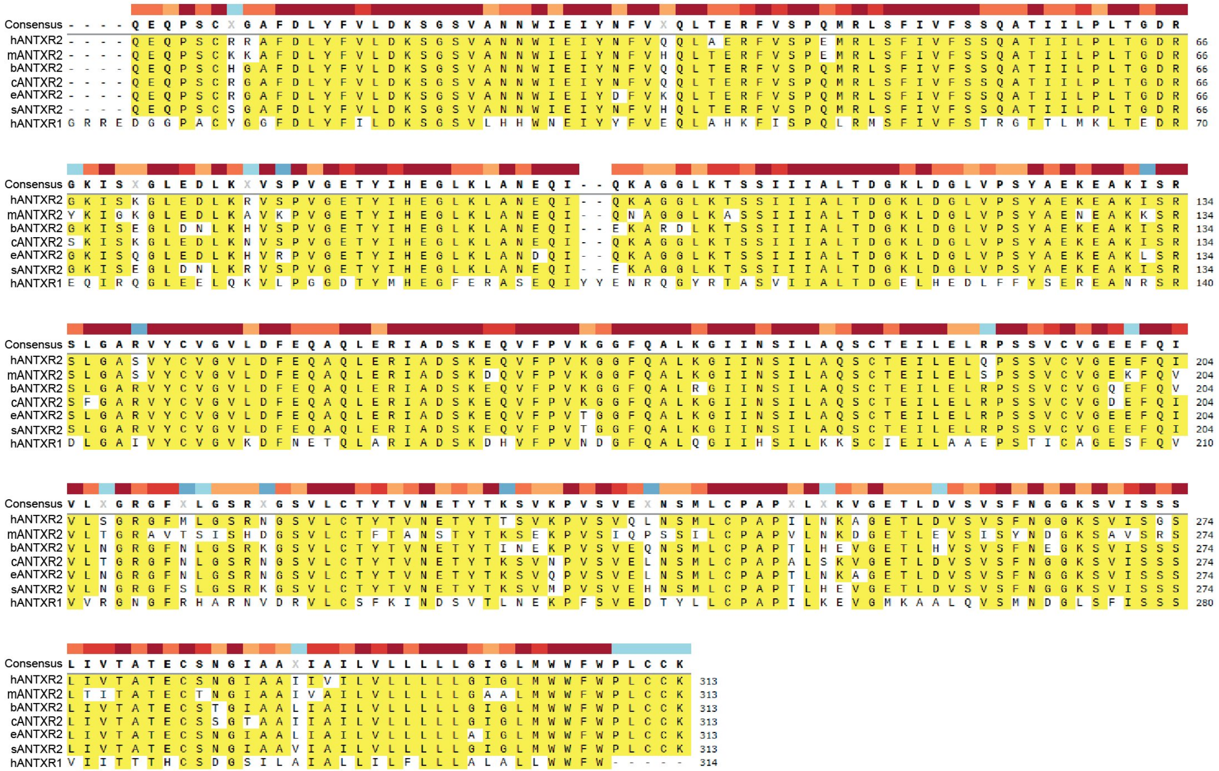

**Sup. Fig. 7.**

**(A)** Sequence alignments of the extracellular and transmembrane domains of ANTXR2 of human, mouse, bovine, canine, equine, swine and human ANTXR1.

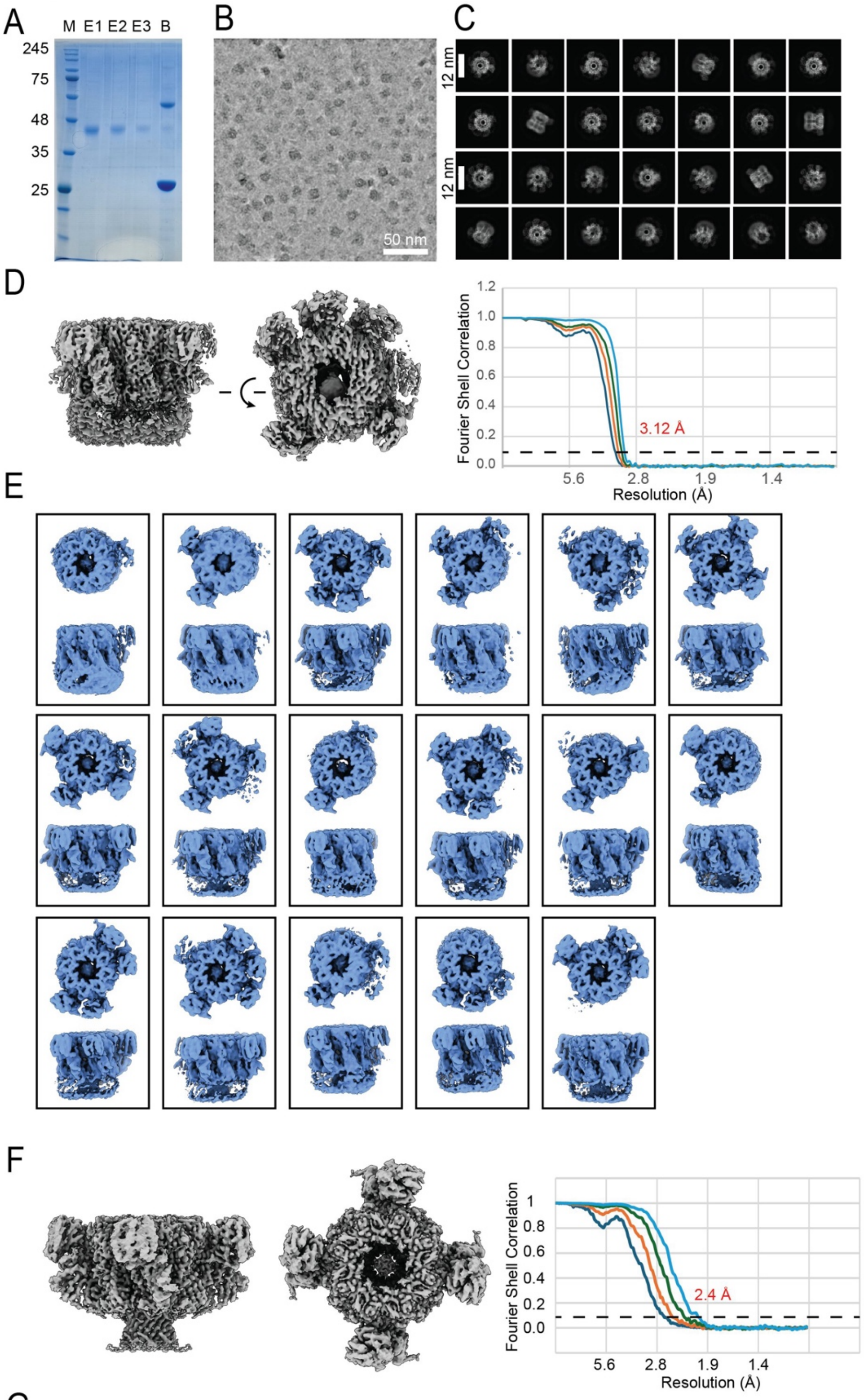


G

EER: 40,082 movies
0.56 Å, 300kV, 40 $e^-/Å^2$
TIF: Convert to 43 frames 0.93 $e^-/Å^2$/frame
Blob pick all micrographs 6,642,714 particles
2D Classification rounds
1,663,758 particles
Ab Initio Reconstruction C1 187,800
3D Classification rounds
Homo Refine 3.12 Å resolution C8
Refinements 2 - 2.8 Å
Local refinements < 3 Å

**Sup. Fig. 8.**

**(A)** FLAG peptide elution fractions (E1-3) of the purification of ANTXR2 from anti-FLAG magnetic beads (B).

**(B)** Cryo-EM micrograph (denoised) of pre-formed NetF pores in DDM:CHS incubated with purified ANTXR2. Scale bar 50nm.

**(C)** Cryo-EM 2D classes of NetF pores with bound ANTXR2.

**(D)** High resolution consensus map of NetF bound to ANTXR2 (no symmetry applied – C1) and FSC plot.

**(E)** 3D classification of the NetF-ANTXR2 complex showing different binding stoichiometries. The consensus model from D was classified in 100 3D classes while filtering to 6 Åresolution to detect the presence or absence of bound ANTXR2. Only significant classes are shown for simplicity.

**(F)** Refinement of a C4 symmetric sub-class of particles and corresponding FSC plot.

**(G)** Brief list of parameters for the acquisition of the NetF:ANTXR2 samples.

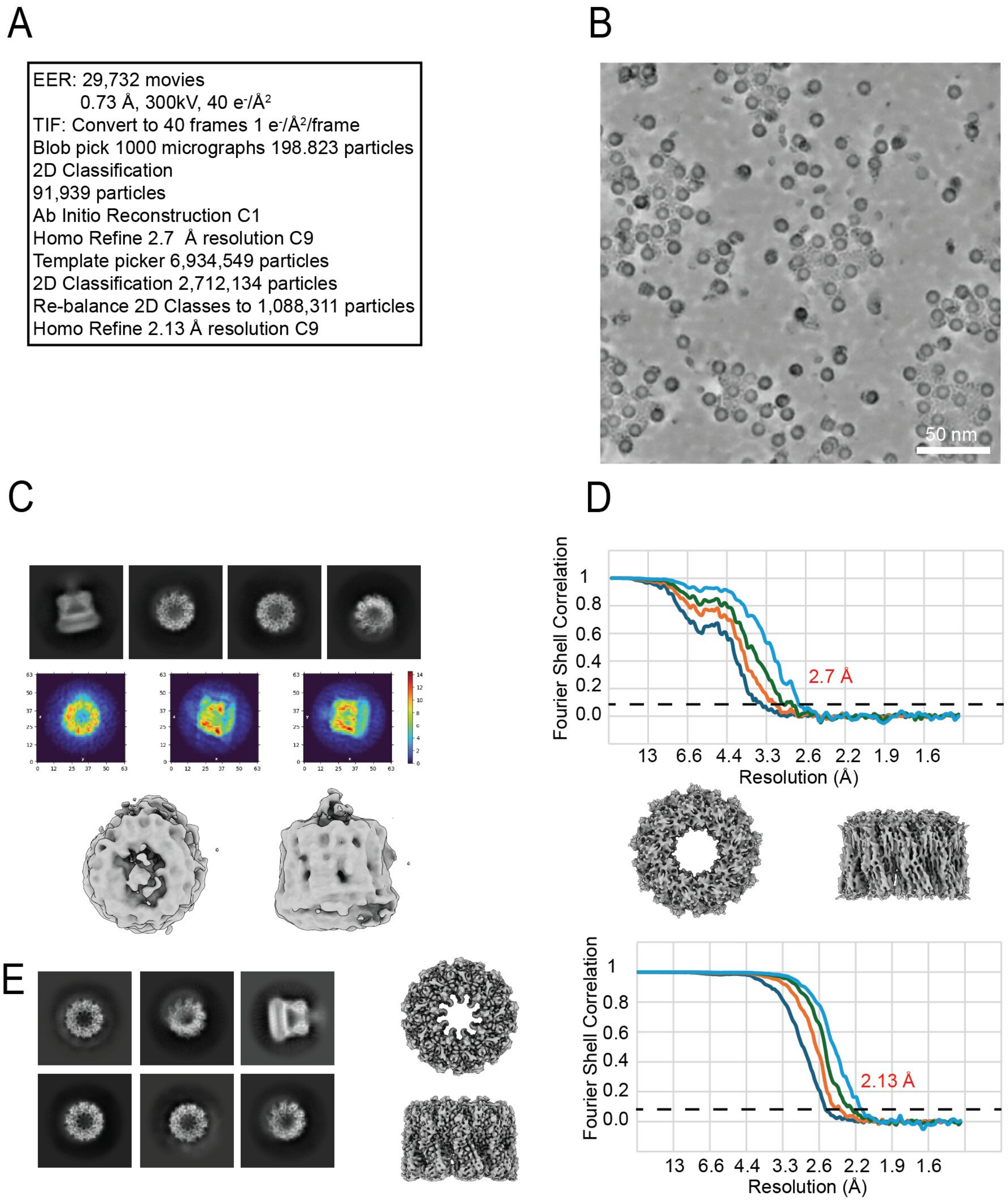


**Sup. Fig. 9.**

**(A)** Acquisition parameters for the NetF pre-pore complex on MSP2N2 nanodiscs in cryo-EM.

**(B)** Representative micrograph of NetF pre-pore assembled on MSP2N2 nanodiscs.

**(C)** Representative 2D classes chosen for initial model generation together with its FSC plot.

**(D)** Representative 2D classes from all the acquired particles.

**(E)** Final NetF pre-pore map together with its FSC plot indicating average resolution obtained.

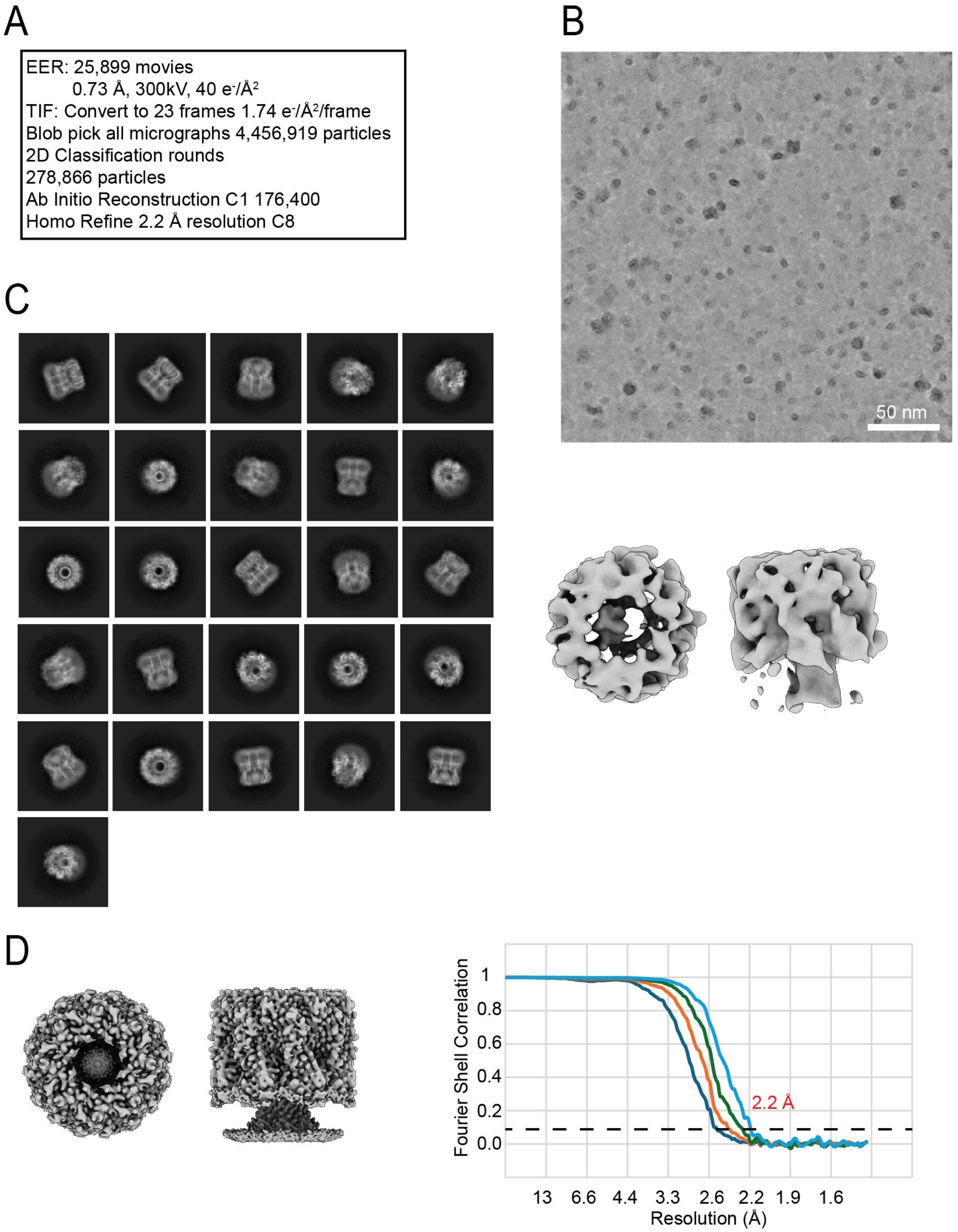


**Sup. Fig. 10.**

**(A)** Acquisition parameters for the NetF pore complex solubilized in DDM:CHS in cryo-EM.

**(B)** Representative micrograph of NetF pore solubilized in DDM:CHS.

**(C)** Representative 2D classes from the Cryosparc pipeline for NetF pore together with the initial model generated.

**(D)** Final NetF pore map together with its FSC plot indicating the average obtained resolution.

| pre-pore | | | | |
|---|---|---|---|---|
| | protomer A | | protomer B | |
| **Number of residues:** | | | | |
| · interface | 44 | 16.50% | 53 | 19.90% |
| · surface | 238 | 89.50% | 237 | 89.10% |
| · total | 266 | 100% | 266 | 100% |
| **Solvent-accessible area, Å** | | | | |
| · interface | 1407.4 | 10.60% | 1431.6 | 10.80% |
| · total | 13313.1 | 100.00% | 13308.5 | 100.00% |
| **Hydrogen bonds** | 25 | | | |
| **Salt bridges** | 7 | | | |
| **pore** | | | | |
| | protomer A | | protomer B | |
| **Number of residues:** | | | | |
| · interface | 59 | 21.90% | 70 | 25.90% |
| · surface | 254 | 94.10% | 253 | 93.70% |
| · total | 270 | 100% | 270 | 100% |
| **Solvent-accessible area, Å** | | | | |
| · interface | 2169.4 | 13.50% | 2162.6 | 13.40% |
| · total | 16128.9 | 100.00% | 16138.2 | 100.00% |
| **Hydrogen bonds** | 40 | | | |
| **Salt bridges** | 7 | | | |
| **NetF - ANTXR2** | | | | |
| | NetF protomer | | ANTXR2 protomer | |
| **Number of residues:** | | | | |
| · interface | 45 | 16.70% | 49 | 17.50% |
| · surface | 252 | 93.30% | 240 | 85.70% |
| · total | 270 | 100% | 280 | 100% |
| **Solvent-accessible area, Å** | | | | |
| · interface | 1473 | 9.10% | 1431.7 | 11.50% |
| · total | 16159.4 | 100.00% | 12495 | 100.00% |
| **Hydrogen bonds** | 12 | | | |
| **Salt bridges** | 6 | | | |

**Table S1**

Summary of the interface residues and areas between two consecutive protomers as calculated by PDBePISA. Upon pore formation the interface is increased from 16-20% to 21-25% mainly by the formation of the extended transmembrane β-barrel.

Summary of the interface residues and areas between a NetF protomer and a ANTXR2 protomer as calculated by PDBePISA. 17% of the residues from each protomer are located in the interface which spans the length of both proteins.

| pre-pore | | | pore | | |
|---|---|---|---|---|---|
| protomer A | protomer B | distance (Å) | protomer A | protomer B | distance (Å) |
| **Hydrogen bonds** | | | **Hydrogen bonds** | | |
| THR 27 | SER 51 | 3.48 | VAL 25 | ASN 50 | 2.85 |
| THR 27 | ASN 52 | 3.45 | GLU 286 | ASN 52 | 3.49 |
| SER 29 | ASN 53 | 3.43 | SER 29 | ASN 53 | 3.39 |
| ASP 30 | LYS 99 | 3.25 | ASP 30 | LYS 99 | 3.08 |
| GLU 150 | THR 108 | 2.71 | GLY 219 | ASN 107 | 2.78 |
| SER 29 | ASN 125 | 3.10 | ASP 153 | GLU 110 | 2.70 |
| LYS 210 | LYS 155 | 2.48 | GLN 151 | VAL 112 | 2.94 |
| SER 217 | LYS 159 | 3.46 | GLN 148 | ARG 113 | 3.66 |
| GLU 33 | ILE 163 | 3.07 | TYR 149 | VAL 114 | 2.97 |
| GLU 208 | ASN 182 | 2.83 | VAL 147 | GLU 116 | 2.79 |
| ASP 30 | LYS 229 | 3.01 | ARG 145 | MET 118 | 2.89 |
| THR 27 | ASN 50 | 2.95 | VAL 143 | TYR 120 | 3.04 |
| SER 28 | ASN 52 | 2.65 | PHE 141 | ILE 122 | 2.86 |
| LYS 61 | GLU 102 | 3.40 | ALA 139 | GLY 124 | 2.83 |
| GLU 150 | ASN 107 | 2.97 | ALA 137 | VAL 126 | 2.89 |
| THR 27 | ILE 122 | 3.85 | VAL 135 | VAL 128 | 2.89 |
| SER 44 | GLY 123 | 2.42 | SER 133 | LYS 131 | 3.82 |
| THR 27 | ASN 125 | 3.03 | LYS 176 | ARG 145 | 3.75 |
| LYS 61 | VAL 126 | 3.73 | LYS 210 | LYS 155 | 2.99 |
| LYS 38 | ASP 161 | 3.00 | GLU 33 | ILE 163 | 3.63 |
| LYS 38 | ASP 161 | 2.90 | ASP 211 | LYS 174 | 2.90 |
| LYS 38 | GLY 162 | 2.86 | GLU 208 | ASN 182 | 3.21 |
| LYS 210 | PHE 172 | 3.19 | ASP 30 | LYS 229 | 2.91 |
| LYS 210 | ASN 173 | 2.84 | THR 27 | ASP 48 | 3.87 |
| LYS 210 | GLN 181 | 3.64 | THR 27 | ASN 50 | 3.32 |
| **Salt bridges pre-pore** | | | SER 28 | ASN 52 | 2.97 |
| ASP 30 | LYS 99 | 3.25 | ASP 153 | GLU 110 | 2.93 |
| GLU 33 | ARG 164 | 3.30 | GLN 151 | VAL 112 | 2.70 |
| GLU 33 | ARG 164 | 3.69 | TYR 149 | VAL 114 | 3.08 |
| ASP 30 | LYS 229 | 3.01 | VAL 147 | GLU 116 | 2.69 |
| LYS 61 | GLU 102 | 3.40 | TYR 149 | GLU 116 | 2.75 |
| LYS 38 | ASP 161 | 2.90 | ARG 145 | MET 118 | 2.97 |
| LYS 38 | ASP 161 | 3.96 | VAL 143 | TYR 120 | 3.03 |
| **Salt bridges pore** | | | PHE 141 | ILE 122 | 2.90 |
| ASP 30 | LYS 99 | 3.08 | ALA 139 | GLY 124 | 2.76 |
| ASP 211 | LYS 174 | 2.90 | ALA 137 | VAL 126 | 2.99 |
| ASP 30 | LYS 229 | 3.96 | VAL 135 | VAL 128 | 3.23 |
| ASP 30 | LYS 229 | 2.91 | LYS 38 | ASP 161 | 3.41 |
| LYS 38 | ASP 161 | 3.04 | LYS 38 | ASP 161 | 3.04 |
| LYS 38 | ASP 161 | 3.62 | LYS 38 | GLY 162 | 2.99 |
| LYS 210 | ASP 180 | 3.20 | | | |

**Table S2**

Residues identified by the PDBePISA algorithm as part of either hydrogen bonds or salt bridges in the interface between consecutive protomers of NetF in pore and pre-pore conformations.

| NetF - ANTXR2 | | |
|---|---|---|
| **NetF protomer** | **distance (Å)** | **ANTXR2 protomer** |
| **Hydrogen bonds** | | |
| ARG 34 | 3.71 | GLU 161 |
| TYR 74 | 2.74 | GLU 200 |
| ARG 264 | 2.76 | GLU 190 |
| ARG 264 | 2.81 | GLU 190 |
| TYR 270 | 2.73 | LEU 241 |
| TYR 273 | 3.43 | GLU 2 |
| ASN 86 | 3.64 | GLN 1 |
| TYR 289 | 3.49 | ILE 176 |
| TYR 273 | 3.12 | SER 179 |
| TYR 289 | 2.82 | SER 179 |
| TYR 271 | 2.88 | ARG 209 |
| GLN 247 | 2.82 | ASN 242 |
| **Salt bridges** | | |
| ARG 34 | 3.71 | GLU 161 |
| ARG 264 | 2.76 | GLU 190 |
| ARG 264 | 2.96 | GLU 190 |
| ARG 264 | 2.81 | GLU 190 |
| ASP 35 | 3.21 | ARG 209 |
| ASP 35 | 3.71 | ARG 209 |

**Table S3**

Residues identified by the PDBePISA algorithm as part of either hydrogen bonds or salt bridges in the interface between a protomer of NetF and ANTXR2.

**Table S4**: Plasmids used in the study

| Purpose | Plasmid name | Source |
|---|---|---|
| Recombinant NetF expression | pET19b_His-NetF | Novagen (EMD Millipore) - cloned by GenScript Biotech |
| Lentivirus production | psPAX2 packaging plasmid | Addgene, #12260 |
| | pMD2-G VSV-G envelope plasmid | Addgene, #12259 |
| Generation of ANTXR2-KO cells | pLentiCRISPR v2 | Addgene, #52961 |
| Expression of constructs in HAP1 cells | pCMV-PuroR | Modified from pcDNA4/TO (Invitrogen) |
| Recombinant ANTXR2 expression | ANTXR2_c1 | Cloned in Addgene, #113900 |
| Expression of ANTXR2 extracellular domain | ANTXR2FLAGHis6 | Modified from LT727211.1 (GenBank) |
| Nanodisc production | pMSP2N2 | Addgene, #29520 |

**Table S5**: Primers and sgRNAs used in the study (overhangs in lower case)

| Purpose | Sequence (5' - 3') | orientation |
|---|---|---|
| ANTXR2 sgRNA1 | caccgAATTTCGTACAGCAACTTG | Forward |
| | aaacCAAGTTGCTGTACGAAATTc | Reverse |
| ANTXR2 sgRNA2 | caccgTTTGATCTCTACTTCGTCC | Forward |
| | aaacGGACGAAGTAGAGATCAAAc | Reverse |
| TIDE analysis for ANTXR2 sgRNA1 (exon 2) | GTGGTTTGCATTTCCTGCGA | Forward |
| | GTCTGAGCCAAGTTTCTGGGA | Reverse |
| TIDE analysis for ANTXR2 sgRNA2 (exon 1) | GCTTCCACTGGGATTCGTCA | Forward |
| | CAAGAGGCCACACTCCCATT | Reverse |
| Amplification of pCMV-puroR backbone to insert hANTXR2 and hANTXR1 | CATCATCACCACCATCACTGAG | Forward |
| | GGTGGCGGATCCTCTAGAGT | Reverse |
| Amplification of hANTXR2 from pRRLSIN.cPPT.PGK-ANTXR2.WPRE to | actctagaggatccgccaccATGGTGGCGGAGCGGTCC | Forward |
| | cagtgatggtggtgatgatgCTGAGATGGAACTCGGGAGAAGTT | Reverse |

| | | |
|---|---|---|
| insert into pCMV-puroR | | |
| Amplification of hANTXR1 from piREShyg2-ANTXR1-HA-137 to insert into pCMV-puroR | actctagaggatccgccaccATGGCCACGGCGGAGCG | Forward |
| | tcagtgatggtggtgatgatgATGGACAGAAGGTCTTGGAGGAGGTCTTG | Reverse |
| Amplification of pCMV-hANTXR2-puroR backbone to insert $GSSS_3$-mNeonGreen | TGAGTCGACAATCAACCTCTGG | Forward |
| | ccgccgctaccgccaccgccCTGAGATGGAACTCGGGAGA | Reverse |
| Amplification of pCMV-hANTXR1-puroR backbone to insert $GSSS_3$-mNeonGreen | TGAGTCGACAATCAACCTCTGG | Forward |
| | ccgccgctaccgccaccgccgctaccgccaccgccGACAGAAGGTCTTGGAGGAGG | Reverse |
| Amplification of $GSSS_3$-mNeonGreen to insert into pCMV-hANTXR2-puroR / pCMV-hANTXR1-puroR | gtagcggcggtggcggtagtggcggtggcggtagtGTGAGCAAGGGCGAGGAG | Forward |
| | aatccagaggttgattgtcgactcaCTTGTACAGCTCGTCCATGC | Reverse |
| Amplification of pCMV-hANTXR2_mNeonGreen-puroR backbone to insert mANTXR2, cANTXR2, eANTXR2 and bANTXR2 | GGCCGGTGCATAAACTTCTCC | Forward |
| | GGTGGCGGATCCTCTAGAGT | Reverse |
| Amplification of mANTXR2 from pcDNA3-ANTXR2_V5 to insert into linearized pCMV-hANTXR2_mNeonGreen-puroR | actctagaggatccgccaccATGGTGGCCGGTCGGTC | Forward |
| | gagaagtttatgcaccggccCTCATCACCTTCCTGGGGTCT | Reverse |
| Amplification of cANTXR2 from pcDNA3.1-cANTXR2 to insert into linearized pCMV-hANTXR2_mNeonGreen-puroR | actctagaggatccgccaccATGCTGGCGGGGCGGAC | Forward |
| | gagaagtttatgcaccggccCTCATCACCTTCCTGGGGTCT | Reverse |
| Amplification of eANTXR2, from pcDNA3.1-eANTXR2, and bANTXR2 from pcDNA3.1-bANTXR2 to insert into linearized pCMV-hANTXR2_mNeonGreen-puroR | actctagaggatccgccaccATGGTGGCGGGGCTGTCC | Forward |
| | gagaagtttatgcaccggccCTCATCACCTTCCTGGGGTCT | Reverse |

| | | |
|---|---|---|
| Removal of Ig domain from pCMV-hANTXR2_mNeonGreen-PuroR | TCTAACGGGATCGCAGCCATC | Forward |
| | AGAGTTTATAATACCATTTAGTGCTTGA | Reverse |
| Removal of vWA domain from pCMV-hANTXR2_mNeonGreen-PuroR | ATACTAGCTCAGTCATGTACTGA | Forward |
| | AAAGGCTCTTCTGCAGGA | Reverse |
| Removal of Ig domain from pCMV-mANTXR2_mNeonGreen-PuroR | ACCAATGGGATTGCAGCC | Forward |
| | AGAGTTGATGATGCCTTTGAGA | Reverse |
| Amplification of hANTXR2 Ig domain from pCMV-hANTXR2_mNeonGreen-PuroR to insert into pCMV-mANTXR2ΔIg_mNeonGreen-PuroR | aaaggcatcatcaactctATACTAGCTCAGTCATGTACTG | Forward |
| | atggctgcaatcccattggtACATTCTGTGGCTGTGAC | Reverse |
| Amplification of mANTXR2 Ig domain from pCMV-mANTXR2_mNeonGreen-PuroR to insert into pCMV-hANTXR2ΔIg_mNeonGreen-PuroR | atggtattataaactctATATTAGCTCGATCATGTACTGA | Forward |
| | atggctgcgatcccgttagaACATTCTGTGGCTGTGATTG | Reverse |
| ANTXR2_AgeI_fw | GTTGCGTAGCTGAAACCGGTATGGTGGCGGAGCGGTCC | Forward |
| ANTXR2_KpnI_rv | AACAGCACCTCAAGGGTACCAAGGGGCCAAAACCACCACA | Reverse |
| Amplification of extANTXR2_insert | ACCTGCCCACCACTGCCGGTCAGGAGCAGCCCTCCTGC | Forward |
| | AAATAAAGATTCTCTGTACGCCCGTTAGAACATTCTGTGGCTG | Reverse |
| Amplification of ANTXR2FLAGHis6_Vector | CGTACAGAGAATCTTTATTTCCAAGGAGGTGC | Forward |
| | ACCGGCAGTGGTGGGCAG | Reverse |

**Table S6**: Amino acid sequences of constructs used in the study (Ig domains are highlighted in yellow).

| |
|---|
| **6xHis-NetF:**<br>MVHHHHHHNSFPESIINSKGKQAEVYTSSDASERDGIKTSLSASFIEDPNSNNLTALVSLKGFIPSGLIKTGTYYSANMYWPSKYNINIETTDEKNNVKILESIPSNTIETVRVTESMGYSIGGNVSVSKKSSSVGANAGFNVQRSVQYEQPDFKTIQKSDGIRKASWNIVFNKTKDGYDQNSYHALYGNQLFMKSRLHNTGAKNLVEDKDLSPLISGGFTPNMVIALKAPKGTKKSMINLNYNLYQDLYTLEWYKTQWWGENRVAKEPYYTYQTYELDWENHTVEFIY |

**Human ANTXR2:**
MVAERSPARSPGSWLFPGLWLLVLSGPGGLLRAQEQPSCRRAFDLYFVLDKSGSVANNWIEI
YNFVQQLAERFVSPEMRLSFIVFSSQATIILPLTGDRGKISKGLEDLKRVSPVGETYIHEGLKLA
NEQIQKAGGLKTSSIIIALTDGKLDGLVPSYAEKEAKISRSLGASVYCVGVLDFEQAQLERIADS
KEQVFPVKGGFQALKGIINSILAQSCTEILELQPSSVCVGEEFQIVLSGRGFMLGSRNGSVLCT
YTVNETYTTSVKPVSVQLNSMLCPAPILNKAGETLDVSVSFNGGKSVISGSLIVTATECSNGIA
AIIVILVLLLLLGIGLMWWFWPLCCKVVIKDPPPPPAPAPKEEEEEPLPTKKWPTVDASYYGGR
GVGGIKRMEVRWGDKGSTEEGARLEKAKNAVVKIPEETEEPIRPRPPRPKPTHQPPQTKWY
TPIKGRLDALWALLRRQYDRVSLMRPQEGDEVCIWECIEKELTA

**Human ANTXR1:**
MATAERRALGIGFQWLSLATLVLICAGQGGRREDGGPACYGGFDLYFILDKSGSVLHHWNEIY
YFVEQLAHKFISPQLRMSFIVFSTRGTTLMKLTEDREQIRQGLEELQKVLPGGDTYMHEGFER
ASEQIYYENRQGYRTASVIIALTDGELHEDLFFYSEREANRSRDLGAIVYCVGVKDFNETQLAR
IADSKDHVFPVNDGFQALQGIIHSILKKSCIEILAAEPSTICAGESFQVVVRGNGFRHARNVDR
VLCSFKINDSVTLNEKPFSVEDTYLLCPAPILKEVGMKAALQVSMNDGLSFISSSVIITTTHCSD
GSILAIALLILFLLLALALLWWFWPLCCTVIIKEVPPPPAEESEEEDDDGLPKKKWPTVDASYYG
GRGVGGIKRMEVRWGEKGSTEEGAKLEKAKNARVKMPEQEYEFPEPRNLNNNMRRPSSPR
KWYSPIKGKLDALWVLLRKGYDRVSVMRPQPGDTGRCINFTRVKNNQPAKYPLNNAYHTSS
PPPAPIYTPPPPAPHCPPPPPSAPTPPIPSPPSTLPPPPQAPPPNRAPPPSRPPPRPSV

**Mouse ANTXR2:**
MVAGRSRARSPGSWLFPGLWLLAVGGPGSLLQAQEQPSCKKAFDLYFVLDKSGSVANNWIEI
YNFVHQLTERFVSPEMRLSFIVFSSQATIILPLTGDRYKIGKGLEDLKAVKPVGETYIHEGLKLA
NEQIQNAGGLKASSIIIALTDGKLDGLVPSYAENEAKKSRSLGASVYCVGVLDFEQAQLERIAD
SKDQVFPVKGGFQALKGIINSILARSCTEILELSPSSVCVGEKFQVVLTGRAVTSISHDGSVLCT
FTANSTYTKSEKPVSIQPSSILCPAPVLNKDGETLEVSISYNDGKSAVSRSLTITATECTNGIAAI
VAILVLLLLLGAALMWWFWPLCCKVVIKDPPPPPSAPMEEEEEEDPLPNKKWPTVDASYYGGR
GVGGIKRMEVRWGDKGSTEEGARLEKAKNAVVMVPEEEIPIPSRPPRPRPTHQAPQTKWYT
PIKGRLDALWALIMKQYDRVSLMRPQEGDEGRCINFSRVPSQ

**Dog ANTXR2:**
MLAGRTLARSPGSRPVPGLWPPLLPLLLLPLLRAPGGPVSAQEQPSCRGAFDLYFVLDNSGS
VANNWIEIYNFVQQLTERFVSPQMRLSFIVFSSQATIILPLTGDRSKISKGLEDLKNVSPVGETYI
HEGLKLANEQIQKAGGLKTSSIIIALTDGKLDGLVPSYAEKEAKISRSFGARVYCVGVLDFEQA
QLERIADSKEQVFPVKGGFQALKGIINSILAQSCTEILELRPSSVCVGDEFQIVLTGRGFNLGSR
NGSVLCTYTVNETYTKSVNPVSVELNSMLCPAPALSKVGETLDVSVSFNGGKSVISSSLIVTAT
ECSSGTAAIIAILVLLLLLGIGLMWWFWPLCCKVVIKDPPPPPPPAPKQEEEEEPLPTKKWPTVD
ASYYGGRGVGGIKRMEVRWGDKGSTEEGARLEKAKNAVVKIPEEVEEPVRPRPPRPKPTYQ
PPQTKWYTPIKGRLDALWALLRRQYDRVSLMRPQEGDEGRCINFSRVPSQ

**Horse ANTXR2:**
MVAGLSLARSPGSWLVPGLWLLVLSGPGGLVSAQEQPSCRGAFDLYFVLDKSGSVANNWIEI
YDFVKQLTERFVSPQMRLSFIVFSSQATIILPLTGDRGKISQGLEDLKHVRPVGETYIHEGLKLA
NDQIQKAGGLKTSSIIIALTDGKLDGLVPSYAEKEAKLSRSLGARVYCVGVLDFEQAQLERIAD
SKEQVFPVTGGFQALKGIINSILAQSCTEILELRPSSVCVGEEFQIVLNGRGFNLGSRNGSVLC
TYTVNETYTKSVQPVSVELNSMLCPAPTLNKAGETLDVSVSFNGGKSVISSSLIVTATECSNGI
AALIAILVLLLLLAIGLMWWFWPLCCKVVIKDPPPPPPPAPKEEEEEPLPTKKWPTVDASYYGG
RGVGGIKRMEVRWGDKGSTEEGARLEKAKNAVVTIPEETEEPVRPRPPRPKPTHQPPQTKW
YTPIKGRLDALWALLRRQYDRVSLMRPQEGDEGRCINFSRVPSQ

| |
|---|
| **Cow ANTXR2:**<br>MVAGLSPARCPGRWLVPGLWLLALSGPGGLLSAQEQPSCHGAFDLYFVLDKSGSVANNWIEI<br>YNFVQQLTERFVSPQMRLSFIVFSSQATIILPLTGDRGKISEGLDNLKHVSPVGETYIHEGLKLA<br>NEQIEKARDLKTSSIIIALTDGKLDGLVPSYAEKEAKISRSLGARVYCVGVLDFEQAQLERIADS<br>KEQVFPVKGGFQALRGIINSILAQSCTEILELRPSSVCVGQEFQVVLNGRGFNLGSRKGSVLC<br>TYTVNETYTINEKPVSVEQNSMLCPAPTLHEVGETLHVSVSFNEGKSVISSSLIVTATECSTGIA<br>ALIAILVLLLLLGIGLMWWFWPLCCKVVIKDPPPPPPSAPKEEEEEPLPTKKWPTVDASYYGG<br>RGVGGIKRMEVRWGDKGSTEEGARLEKAKNAVVTLPEEPEEPVRPRPPPSKPTHQPPQTK<br>WYTPIKGRLDALWALLRRQYDRVSLMRPQEGDEGRCINFSRVPSQ |
| **GSSS$_3$-mNeonGreen:**<br>GGGGSGGGGSGGGGSVSKGEEDNMASLPATHELHIFGSINGVDFDMVGQGTGNPNDGYE<br>ELNLKSTKGDLQFSPWILVPHIGYGFHQYLPYPDGMSPFQAAMVDGSGYQVHRTMQFEDGA<br>SLTVNYRYTYEGSHIKGEAQVKGTGFPADGPVMTNSLTAADWCRSKKTYPNDKTIISTFKWSY<br>TTGNGKRYRSTARTTYTFAKPMAANYLKNQPMYVFRKTELKHSKTELNFKEWQKAFTDVMG<br>MDELYK |
| **extANTXR2-FLAG-His6:**<br>MGTLSAPPCTQRIKWKGLLLTASLLNFWNLPTTAGQEQPSCRRAFDLYFVLDKSGSVANNWI<br>EIYNFVQQLAERFVSPEMRLSFIVFSSQATIILPLTGDRGKISKGLEDLKRVSPVGETYIHEGLK<br>LANEQIQKAGGLKTSSIIIALTDGKLDGLVPSYAEKEAKISRSLGASVYCVGVLDFEQAQLERIA<br>DSKEQVFPVKGGFQALKGIINSILAQSCTEILELQPSSVCVGEEFQIVLSGRGFMLGSRNGSVL<br>CTYTVNETYTTSVKPVSVQLNSMLCPAPILNKAGETLDVSVSFNGGKSVISGSLIVTATECSNG<br>RTENLYFQGGAGARSIEGRIVKDYKDDDDKHHHHHH |

**Dataset 1**

Differential gene expression analysis of U937-R and J3T-R cells compared to their respective wild-type controls (see attached excel file).

This table includes RNA-seq data from U937 and J3T cells, comparing resistant (R) and wild-type (WT) variants in triplicates. The first tab (U937 RNA-seq) presents expression values (TPM) for U937-R and U937-WT cells. The second tab (J3T RNA-seq) provides analogous data for J3T-R and J3T-WT cells. In each tab, columns B–I show expression values from individual samples, columns J–L list mean expression per condition, column L gives $\log_2$ fold change (R vs. WT), and columns M-N report p-values and adjusted p-values (padj) from differential expression analysis. Column O includes gene names.
